# G-quadruplexes represent promising new targets to overcome multidrug-resistant fungal infections

**DOI:** 10.1101/2025.03.01.640902

**Authors:** Georgie Middleton, Fuad O Mahamud, Isabelle Storer, Abigail Williams-Gunn, Finn Wostear, Alireza Abdolrasouli, Elaine Barclay, Alice Bradford, Oliver Steward, Silke Schelenz, James McColl, Bertrand Lézé, Norman van Rhijn, Alessandra da Silva Dantas, Takanori Furukawa, Derek Warren, Zoë AE Waller, Stefan Bidula

## Abstract

The growing emergence of antifungal resistance has prompted the identification of novel antifungal targets. G-quadruplexes (G4s), four-stranded secondary structures that form in DNA and RNA, have arisen as a drug target to treat bacterial, viral, and parasitic infections. Here, we provide the first demonstration that G4s form in fungi and represent a promising new target for antifungal development. We found that PhenDC3 and pyridostatin (PDS), ligands that bind to and stabilise G4s, potently inhibited the metabolism of fungal pathogens in different ways. These pathogens included the pan azole-resistant *Aspergillus fumigatus* isolate TR_34_/L98H and *Candida auris*. Notably, PhenDC3 could synergise with amphotericin B, was protective in an *in vivo* model of fungal infection, and was well-tolerated by human cells. However, continuous exposure to PhenDC3 resulted in some cross-resistance to current antifungals and this needs to be explored further. PhenDC3 could also increase the number of RNA G4s in live *A. fumigatus*. Further, we demonstrated the structural flexibility of DNA sequences found in *cyp51A* and *cyp51B*, with these sequences capable of forming duplexes, hairpins, G4s and i-motifs. Finally, PhenDC3, but not PDS, caused duplex DNA structures in *cyp51A* to transition into antiparallel G4 structures potentially associated with PhenDC3’s increased antifungal potency. Taken together, G4s represent an exciting antifungal target, but a more detailed understanding of their biological roles is essential.

## Introduction

Pathogenic fungi represent a threat to humanity and contribute to the loss of almost 4 million lives annually.^1^ This would be an alarming statistic on its own, but fungal infections also devastate critical food crops (e.g., cereals, wheat, rice) and significantly impact biodiversity.^2^ With an increase in the utilisation of immunosuppressive medical interventions, increased incidence of pandemic level infections, and global warming; the frequency of life-threatening fungal infections is likely to rise.^3^ Worryingly, this rise coincides with an emergence of drug resistance, which has prompted the World Health Organization (WHO) to respond and compile a list of priority fungal pathogens.^4^ This increase in drug resistance has been found to be associated with the overuse of both azole pesticides and more recently, dihydroorotate dehydrogenase (DHODH) inhibitors, in agricultural practices.^5–9^ Due to a limited number of effective antifungal agents and the emergence of resistance, novel antifungals with an alternative mechanism of action are highly desired.

G-quadruplexes (G4s) are non-canonical four-stranded secondary structures that form within guanine-rich regions of DNA and RNA.^10^ Rather than Watson-Crick base pairing, four guanines can associate with one another via Hoogsteen hydrogen bonding to form G-tetrads. These G-tetrads are connected by loops of nucleotides and can π-π stack to form the G4 structure, with the number of G-tetrads often proportional to G4 stability.^11^ Alternatively, another non-canonical structure, the i-motif (iM), can form in the cytosine-rich complementary strand through hemi-protonated cytosine-cytosine base pairing, and the formation of G4s and iMs are interdependent.^12,13^

Due to the enrichment of G4s/iMs in regulatory regions of the genome, such as the promoters, untranslated regions, and introns, their roles in regulating fundamental biological processes such as transcription, translation, alternative splicing, and epigenetics have become abundantly apparent.^10,14,15^ G4-ligands, such as PhenDC3 and pyridostatin (PDS), have been designed to specifically stabilise G4s within the genome with negligible interaction to B-DNA.^16,17^ Although much investment has been placed in the therapeutic potential of G4-ligands in the treatment of cancer, there is growing evidence of the antibacterial, antiviral, and antiparasitic possibilities of G4-ligands.^18–20^ However, the antimicrobial potential of targeting fungal G4s has been poorly explored. Here, we provide the first comprehensive study highlighting the formation of G4s in pathogenic fungi and the therapeutic potential of targeting non-canonical nucleic acid structures in both drug-susceptible and drug-resistant critical priority fungal pathogens.

## Results

### The G4-ligands PhenDC3 and PDS display antifungal activity against Aspergillus spp. and Candida spp

Given the increased prevalence of antifungal resistance, identifying drugs with novel mechanisms of action is of utmost importance. To explore the antifungal potential of G4-ligands, we tested the antifungal activity of PhenDC3 and PDS against a panel of *Aspergillus* and *Candida* species.

PhenDC3 inhibited the metabolic activity of a range of *A. fumigatus* isolates. The MIC_90_ for the azole-susceptible *A. fumigatus* clinical isolate 22M7007854 was 1.56 µM (**Fig. 1a**). Surprisingly, PhenDC3 had increased activity against 22M7004177 (MIC_90_ = 0.78 µM) and TR_34_/L98H (MIC_90_ = 0.78 µM); itraconazole and pan azole-resistant isolates, respectively. PhenDC3 also retained its activity against two common *A. fumigatus* laboratory strains, CEA10 (MIC_90_ = 3.13 µM) and A1160+ (MIC_90_ = 3.13 µM). Conversely, PDS had activity against all the isolates tested, but its efficacy was lower compared to PhenDC3 (MIC_90_ = 12.50 µM for 22M7007854, 22M7004177, and TR_34_/L98H; MIC_90_ = 6.25 μM for CEA10 and A1160+; **Fig. 1b**).

**Figure 1:**
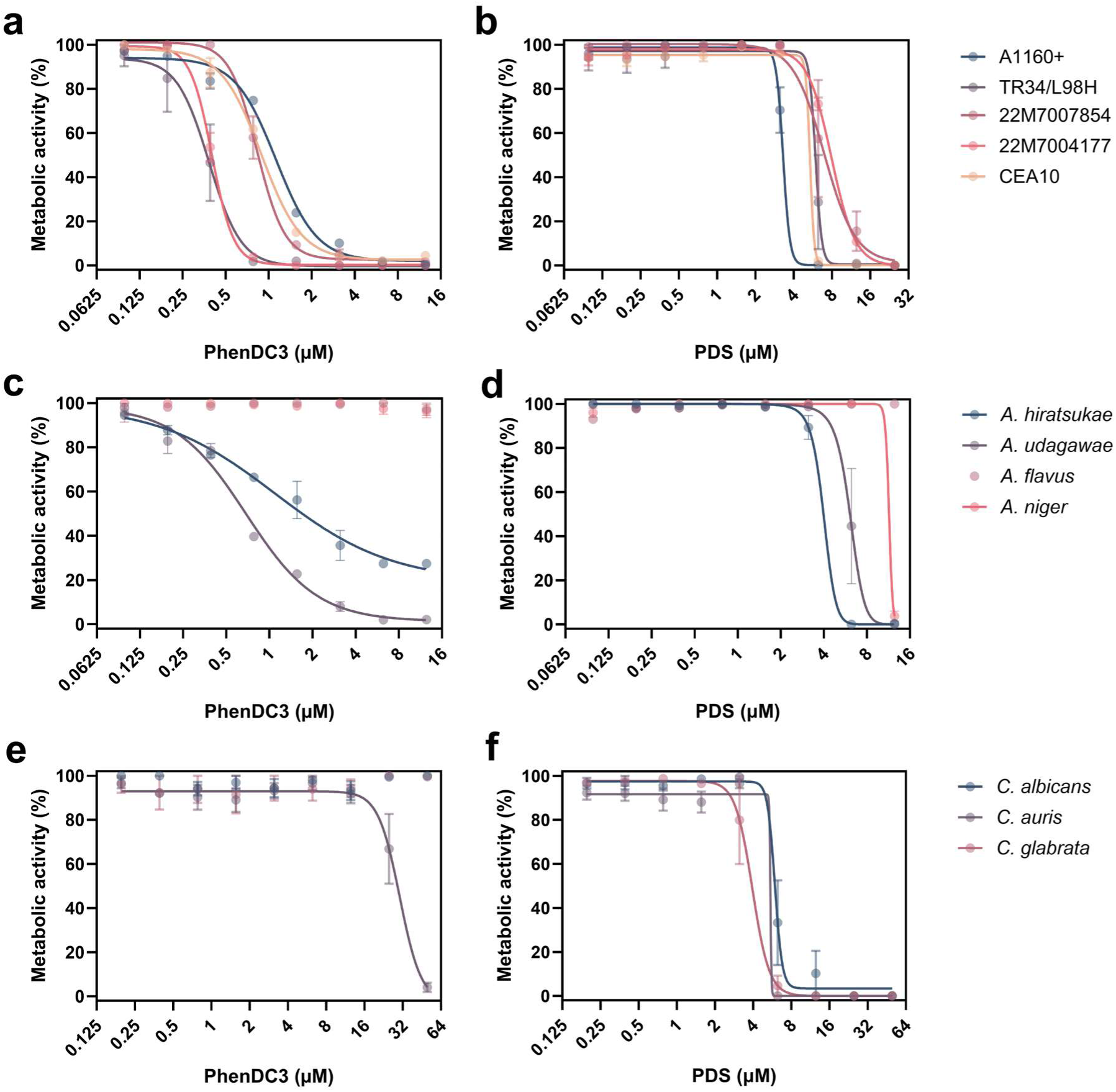
PhenDC3 and PDS inhibit the metabolism and growth of drug susceptible and drug-resistant fungal pathogens. Drug susceptible (22M7007854) and itraconazole-resistant (22M7004177) *A. fumigatus* clinical isolates, laboratory strain A1160+, ‘wildtype’ strain CEA10, and the pan azole-resistant *A. fumigatus* isolate (TR34/L98H) were treated with a concentration range of (**a**) PhenDC3 or (**b**) PDS for 48 h at 37 °C. Non-*fumigatus Aspergillus* strains treated with (**c**) PhenDC3 or (**d**) PDS for 48 h. *Candida* species were treated with the indicated concentrations of (**e**) PhenDC3 and (**f**) PDS for 24 h. Metabolic activity was calculated as percentage of the untreated control. Experiments are in biological triplicate with each data point consisting of the mean of three technical replicates ± SEM.

Next, we investigated the effects of PhenDC3 against non-*Fumigati* and cryptic *Fumigati* species to see if our observations were transferable to related fungi. PhenDC3 appeared to lose activity against the non-*Fumigati* species *A. flavus* and *A. niger*, but it was still inhibitory against the cryptic *Fumigati* species *A. udagawae* (MIC_90_ = 6.25 µM) and *A. hiratsukae* (MIC_90_ = >12.5 µM), which are genetically very similar to *A. fumigatus* (**Fig. 1c**). PDS was also inactive against *A. flavus* but displayed activity against *A. niger* (MIC_90_ = 12.5 μM), *A. hiratsukae* (MIC_90_ = 6.25 μM), and *A. udagawae* (MIC_90_ = 12.5 μM) (**Fig. 1d**).

Finally, we assessed the antifungal activity of both PhenDC3 and PDS against the *Candida* spp. commonly associated with human infections; *C. albicans, C. glabrata*, and *C. auris,* to determine if G4-ligands had activity against fungi more distantly related to *Aspergilli*. Surprisingly, PhenDC3 displayed poor activity against *C.* auris (MIC_50_ = 30.03 μM) and no activity against the other *Candida* spp. tested (**Fig. 1e**). However, PDS was found to be more active against these yeasts compared to filamentous fungi and displayed potent activity (MIC_50_ = 5.48 μM, 3.94 μM, and 5.94 μM for *C. auris, C. glabrata,* and *C. albicans*, respectively; **Fig. 1f**). A full list of potencies can be found in **Supplementary Table 1**.

### PhenDC3 and PDS significantly impact A. fumigatus growth and morphology

To determine if PhenDC3 and PDS were fungicidal or fungistatic, we performed a killing assay on CEA10. Both PhenDC3 and PDS exhibited apparent fungicidal activity in a concentration dependent manner 24 h after treatment, with MIC values of 1.56 µM and 12.5 µM being recorded for PhenDC3 and PDS, respectively (**Fig. 2a**). Colonies appeared 48 h following treatment with PhenDC3, suggesting a fungistatic effect. However, the phenotype of the colonies recovered 48 h post treatment were smaller and less sporulated than untreated colonies. Additionally, colonies with a similar phenotype were recovered following treatment with the known fungicidal antifungal, amphotericin B **(Fig. 2b)**. To explore how quickly G4-ligands could exert their antifungal effects, we performed a kill-curve analysis. We found that G4-ligands had a fast mechanism of action: MIC concentrations of PhenDC3 and PDS prevented >90% of growth after 24 h of outgrowth following 8 hours and 1 hour of treatment, respectively (**Fig. 2c, d**).

**Figure 2:**
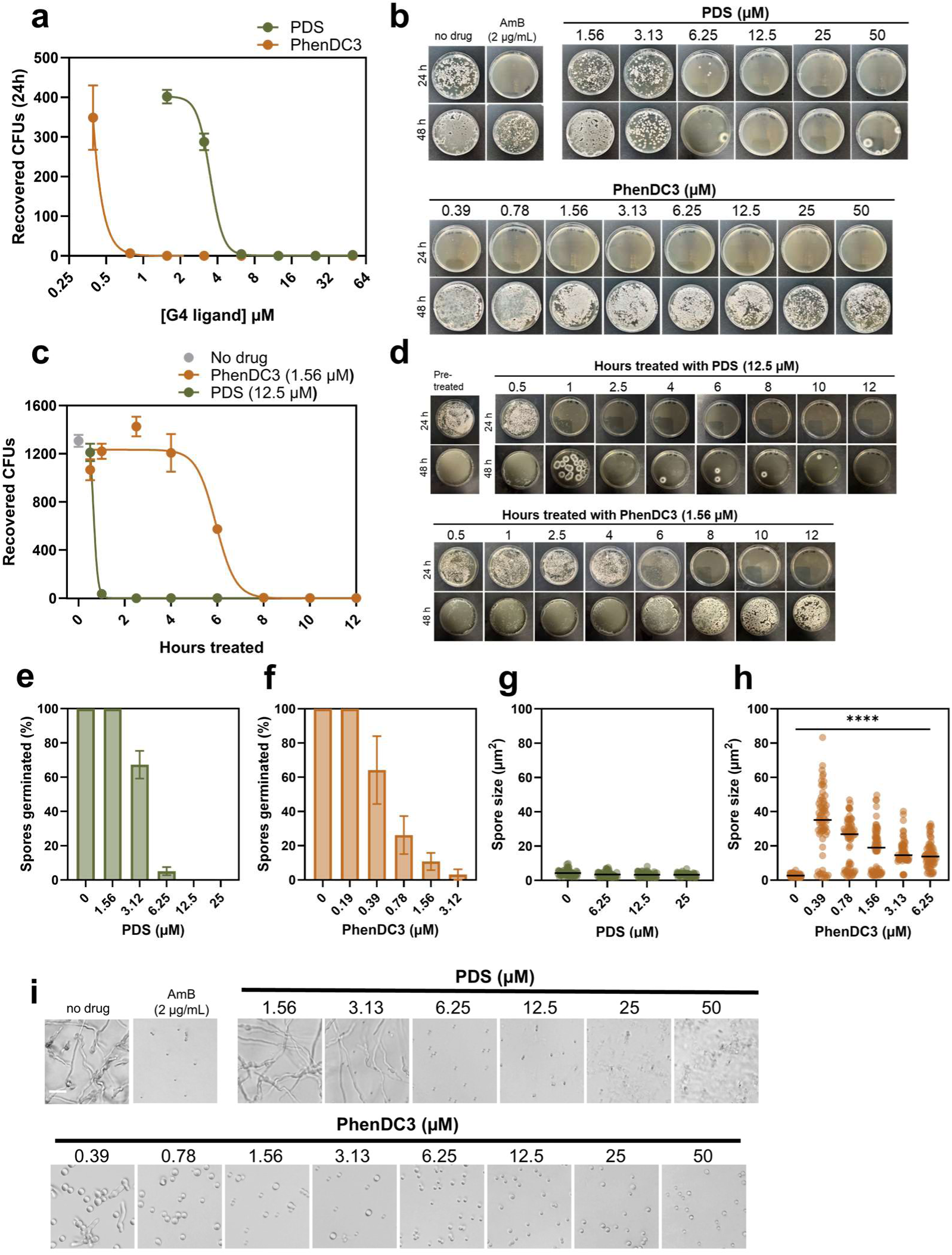
PhenDC3 exhibits fungistatic activity after eight hours of treatment, and PDS exhibits fungicidal activity after just one hour of exposure to *A. fumigatus* conidia. (**a**) Viability counts and (**b**) colony phenotypes of *A. fumigatus* conidia (CEA10) after 24 h of exposure to the indicated range of PDS and PhenDC3. SDA plates were incubated for 24 h and 48 h hours post exposure. Counts are the mean of three technical replicates. (**c**) Viability counts and (**d**) colony phenotypes of *A. fumigatus* conidia (CEA10) after the indicated time of exposure MIC90 concentrations of PDS and PhenDC3. SDA plates were incubated for 24 h and 48 h hours post exposure. Counts are the mean of three technical replicates. The percentage of conidia which germinated after 24 h of exposure at 37°C to (**e**) PDS or (**f**) PhenDC3. The mean ± SEM of three technical replicates is shown. The area of conidia which did not germinate were measured using Fiji 1.53t (n=60) after 24 h exposure to (**g**) PDS or (**h**) PhenDC3. Statistical significance was determined by a Kruskal-Wallis test with Dunn’s multiple comparisons correction (H (5) = 174.5, *p*<0.0001). untreated vs. each PhenDC3 treatment concentration were significantly different (*p*<0.0001, (****)). (**i**) Images of *A. fumigatus* conidia treated for 24 h with the indicated concentration of G4-ligand. Scale bar: 20 μm.

The rapid mechanism of action of PhenDC3 and PDS was mirrored in their ability to prevent *A. fumigatu*s from germinating, and both compounds could inhibit germination in a concentration-dependent manner (**Fig. 2e, f**). PhenDC3 also had activity against germlings but no activity against mature biofilms at the concentrations tested (**Supplementary** Fig. 1a).

Non-germinated conidia treated with PDS did not swell, which may be attributed to the fast killing exhibited by PDS (**Fig. 2g**). Rather than germinating, conidia treated with PhenDC3 became excessively swollen: the mean untreated spore area was 2.66 μm ± 0.16 SEM, whilst the mean spore area when treated with 0.39 μM PhenDC3 was 34.38 μm ± 2.30 SEM (**Fig. 2h, Supplementary** Fig 1b). However, a small subset of conidia did not appear to swell or germinate. Images of conidia treated for 24 h with PDS and PhenDC3 are found in **Fig. 2i**.

### G4s can form in both the RNA and DNA of A. fumigatus

The frequency of G4s in the *A. fumigatus* reference genome, Af293, was found to be comparable to the G4-frequency observed in humans at around 1.5 per 1000 base pairs when using the default G4Hunter settings (unpublished). This is far higher than our initial estimations as these parameters account for the detection of non-canonical G4s which were filtered out in our previous analyses.^21^ Regardless, the formation of G4s or iMs in *A. fumigatus* RNA and DNA had never been confirmed experimentally.

To determine RNA G4 formation in live fungi, we used QUMA-1, a dynamic dye that fluoresces when it binds to RNA harbouring G4 structures.^22^ In these experiments, Hoechst 33342 was intended to be used to label the fungal nuclei. However, it did not enter live fungi when incubated for a short period of time, and this was used instead to help outline the fungus and enhance visualisation. QUMA-1 staining was diffuse in untreated *A. fumigatus,* with few foci observed (mean 0.35 ± 0.11 SEM foci per 10 µm^2^) (**Fig. 3a**). Notably, PhenDC3 induced an obvious increase in the number of punctate QUMA-1 foci at both 1.56 µM (mean 2.65 ± 0.26 SEM foci per 10 µm^2^) and 12.5 µM (mean 3.09 ± 0.39 SEM foci per 10 µm^2^) **(Fig. 3b**), suggesting that PhenDC3 was influencing G4-formation directly in fungi as anticipated. Fungi treated with 1.56 µM PDS exhibited a phenotype comparative to control but 12.5 µM PDS prevented the swelling of conidia. It was impossible to determine the foci in the 12.5 µM PDS-treated spores because the spores were dead, and the membrane was more permeable to both dyes.

**Figure 3:**
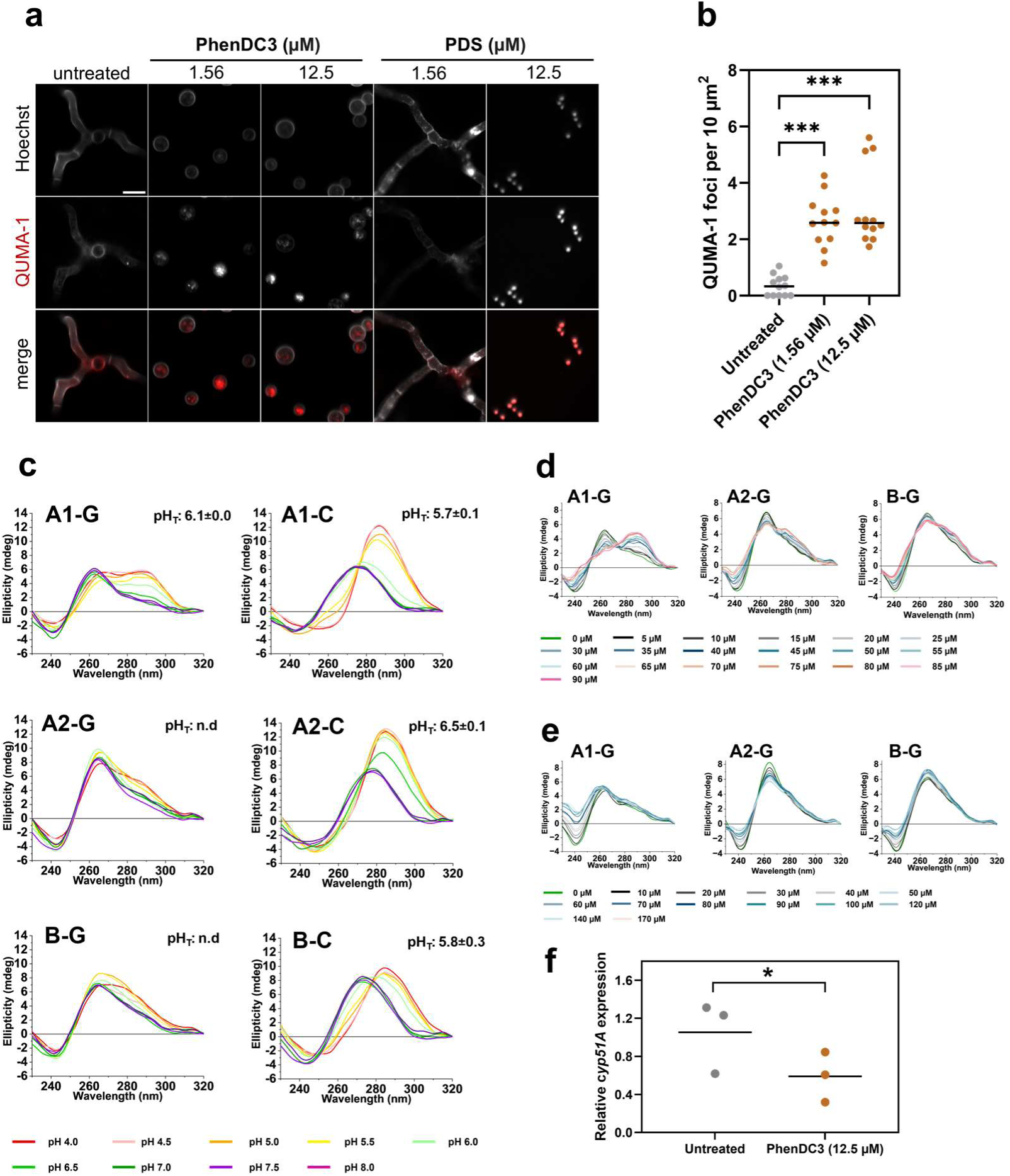
G4s form both within *A. fumigatus* RNA *in vivo*, and *cyp51A* and *cyp51B* gene sequences *in vitro*. (**a**) *A. fumigatus* conidia were treated with the indicated concentration of G4-ligand for 18 h at 37 °C. Treated cells were washed with PBS and incubated with 1 µM Hoechst 33342 and 1 µM QUMA-1 at 37 °C for 1 h to observe RNA G4. Representative images were captured via confocal microscopy using a 100x oil objective. Scale bar: 10 µm. (**b**) The number of QUMA-1 foci per cell per image (n=12) was quantified using Fiji 1.53t and normalised per area to allow for differences in germination. Statistical significance was determined by a Kruskal-Wallis test with Dunn’s multiple comparisons correction (H (2) = 3.47, *p*<0.001). Untreated vs. PhenDC3 (1.56 µM) or (12.5 µM) were significantly different (*p*<0.001, (***)). (**c**) CD spectroscopy of the potential G4-forming oligonucleotide sequences found in the *cyp51A* coding region (A1), 3’-untranslated region (A2), and *cyp51B* coding region (B). Both C-rich and G-rich strands are shown. All spectra were obtained by annealing 10 µM DNA oligonucleotide in a buffer containing 10 mM sodium cacodylate and 100 mM KCl at pH 4 to 8 as indicated. The transitional pH (pHT) was noted when a structural shift was observed and was determined from the inflection point of a Boltzmann sigmoidal. The ellipticity of the c*yp51A* and c*yp51B* oligonucleotide sequences (10 µM) at 295 nm was determined in the presence of (**d**) 0-90 µM PhenDC3 or (**e**) 0-170 µM PDS in 10 mM sodium cacodylate and 100 mM KCl buffer at pH 7.0. Data are representative of the ellipticity at 295 nm observed in two biological replicates. Error bars represent the mean ±SD. (**f**) *cyp51A* expression was quantified by RT-qPCR. *A. fumigatus* conidia were incubated with or without 12.5 μM PhenDC3 for 4 h. RNA was extracted by TRIzol-chloroform. Primers were designed across exon-exon junctions. Cyp51A expression was normalised to TubA. Experiments were performed in biological triplicate. Each data point represents the mean of three technical replicates. Statistical significance was determined by a paired t test (untreated mean 1.05 (± 0.21 SEM), PhenDC3 treated mean 0.59 (± 0.15 SEM). t(4) = 4.938, *p*<0.05, (*)).

Our previous bioinformatic analysis identified G4-forming sequences in *cyp51A* and *cyp51B*, two sterol 14-demethylases critically involved in the synthesis of ergosterol in the fungal cell membrane and the target of azole antifungals.^21^ These sequences were found in the *cyp51A* coding and 3’-untranslated regions (termed A1-G and A2-G, respectively), and the coding region of *cyp51B* (termed B-G). Here, we biophysically characterised these three sequences and their reverse complement (A1-C, A2-C, and B-C) as a proof of concept to demonstrate G4 formation in fungal DNA. We found that these DNA sequences displayed a large amount of structural flexibility depending upon the cations present or the pH and were capable of folding into G4s, iMs, duplexes, and hairpins as determined by circular dichroism (CD), thermal difference spectra (TDS) and UV-melting experiments (**Table 1**).

**Table 1:**
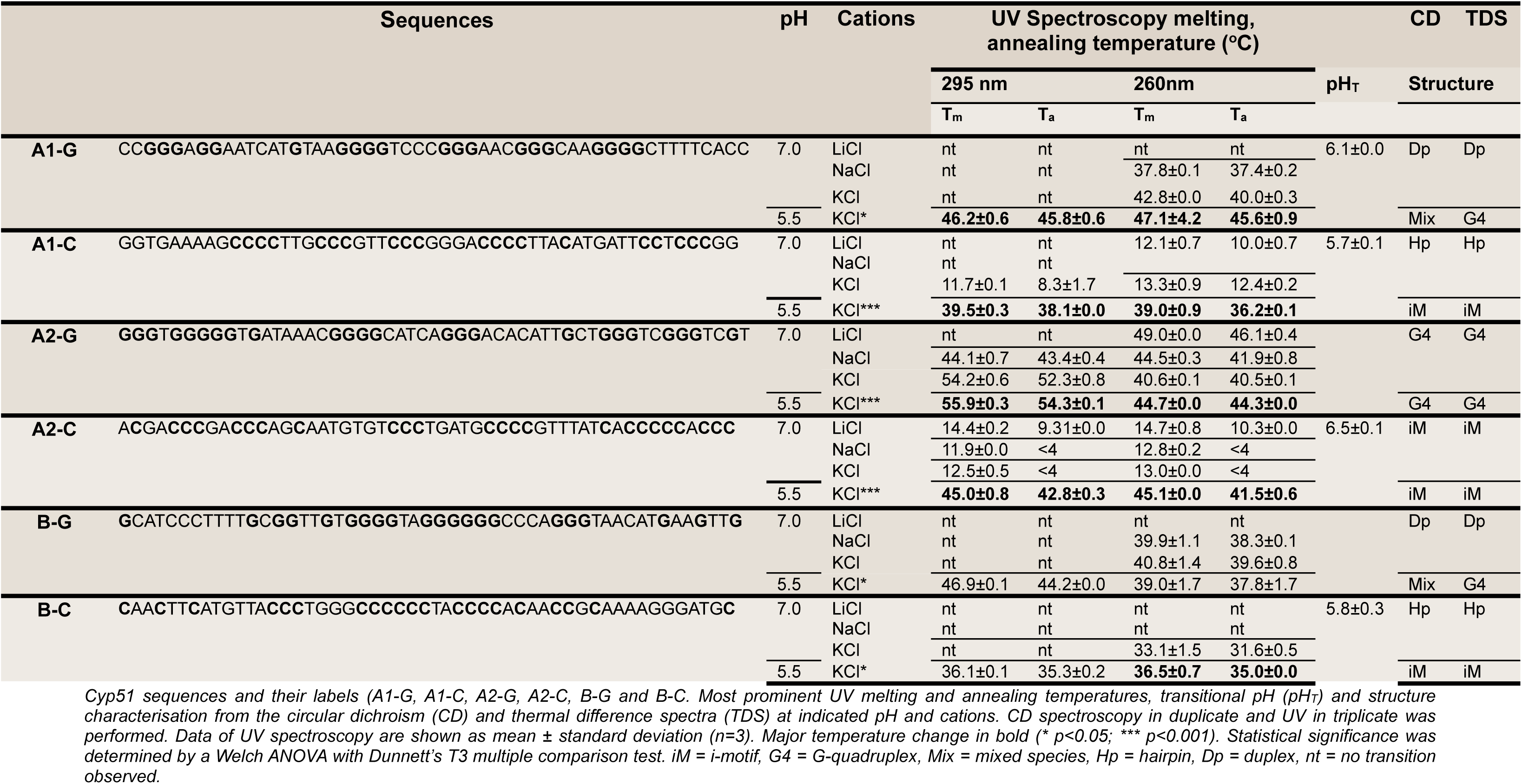
Biophysical characterisation data for the Cyp51 sequences.

At pH 7.0, A1-G and B-G gave CD spectra with a broad positive peak at 260 nm and a negative peak at 245 nm (**Fig. 3c, bottom panel**). This type of spectra could indicate either a parallel G4 or duplex structures. Complementary TDS confirmed duplex DNA conformation, rather than G4, with a notable absence of a negative band at 295 nm.^23^ UV-melting experiments showed transitions only at 260 nm, in-line with duplex structures, and not at 295 nm, which would be expected for G4s. Unexpectedly, as the pH of the samples were lowered, the CD spectra of both A1-G and B-G transitioned into antiparallel G4 structures, as indicated by an increased signal at 295 nm in the CD spectra. This change was consistent with a pH-dependent conformational shift, where the degree of protonation of cytosine bases affects the stability and folding of G4 structures.^24^ These duplex-to-G4 transitions were supported by the UV melting experiments (i.e., increased melting temperature of G4 versus duplex in KCl at pH 5.5) and TDS (positive peaks at 240 nm, 255 nm, and 270 nm, and a negative peak at 295 nm). All supporting data for these and the following biophysical experiments can be found in **Supplementary** Fig. 2-6.

The C-rich sequences A1-C and B-C transitioned between hairpin and iM structures depending on the cationic and pH conditions (**Fig. 3c, top panel)**. CD spectra at pH 7.0 showed positive ellipticity at 272 nm, indicative of hairpin formation. As the pH decreased, the spectra shifted to a maximum at 288 nm, and negative peak at 240 nm, consistent with the formation of an iM structure. The TDS profile for these sequences showed positive peaks at 240 nm and 265 nm with a negative peak at 295 nm, characteristic of iM conformation at pH 5.5. A1-C also exhibited distinct UV melting and annealing profiles at 260 nm at pH 7.0 under different cationic conditions yet only displayed a melting profile at 295 nm in KCl at pH 5.5.

The C-rich sequences A1-C and B-C transitioned between hairpin and iM structures depending on the pH conditions. CD spectra at pH 7.0 showed a positive peak at 272 nm, indicative of hairpin formation. As the pH decreased, this peak shifted to a maximum at 288 nm, and a negative peak at 240 nm, consistent with the formation of an iM structure. The TDS profile for these sequences showed positive peaks at 240 nm and 265 nm with a negative peak at 295 nm, characteristic of iM conformation. A1-C also exhibited distinct UV melting and annealing profiles at 260 nm across various cationic conditions and displayed a melting profile at 295 nm in KCl at pH 5.5 and pH 7.0.

A2-G adopted both hairpin and G4 conformations under physiological-like conditions (**Fig. 3c, middle panel**). This was characterised by a positive peak at 263 nm, with a shoulder at 280 nm, and a negative peak around 240 nm, indicating the co-existence of two structural forms. This result was further supported by UV melting data measured at 260 nm and 295 nm. In KCl at pH 7.0, the melting temperature from the data recorded at 260 nm was 40.6 ±0.1°C, corresponding to the hairpin structure, whereas the melting temperature at 295 nm was significantly higher (54.2 ±0.6°C), indicative of the presence of a more thermodynamically stable G4 structure.

Sequence, A2-C formed iM structures and was significantly more stable at pH 5.5 compared to pH 7.0.

We examined the interactions of PhenDC3 and PDS on the G-rich sequences A1-G, A2-G, and B-G (**Fig. 3d, e**). PhenDC3 was found to cause the duplex structures in A1-G and parallel G4 structures in A2-G to transition to antiparallel G4 conformations. The transition from a duplex structure to an antiparallel G4 in A1-G was particularly striking. Binding of PhenDC3 was found to be cooperative, with a K value of 32.7 ± 3.3 µM and a Hill coefficient of 2.1 ± 0.9 when interacting with PhenDC3 (**Supplementary** Fig. 7). Similarly, A2-G showed a comparable binding affinity, with a K value of 32.5 ± 1.8 µM and a Hill coefficient of 2.8 ± 1.5 for PhenDC3 (no difference between PhenDC3 binding to A1-G or A2-G; p=0.9486). Contrarily, PhenDC3 had limited effect on the duplex structure formed in B-G. Importantly, PDS did not induce any structural alterations in any of the sequences tested. In support of PhenDC3’s influence on these *cyp51A* sequences (A1-G and A2-G), PhenDC3 reduced the expression of *cyp51A* by 44% relative to control (p<0.05; **Fig. 3f**). We were unable to obtain high-quality RNA from the PDS-treated fungi, presumably due to the fast killing by PDS.

### Targeting fungal G4s may have therapeutic potential

The above data demonstrated that PhenDC3 possessed impressive antifungal activity and could directly influence the number and structure of RNA and DNA G4s, respectively. Next, we explored whether PhenDC3 had the potential to be utilised therapeutically and assessed the synergy of G4-ligands with current antifungals, the *in vivo* efficacy of PhenDC3, and the effect of G4-ligands on disease-relevant human cells.

Checkerboard assays performed with CEA10 conidia indicated that PhenDC3 could synergise with amphotericin B (FICI = 0.375), with a combination of 0.39 μM PhenDC3 and 0.125 µg/mL amphotericin B able to fully inhibit germination (**Fig. 4a**). Meaning that 8-fold lower concentrations of amphotericin B and 4-fold lower concentrations of PhenDC3 could be used in combination. PhenDC3 did not synergise with itraconazole or voriconazole.

**Figure 4.**
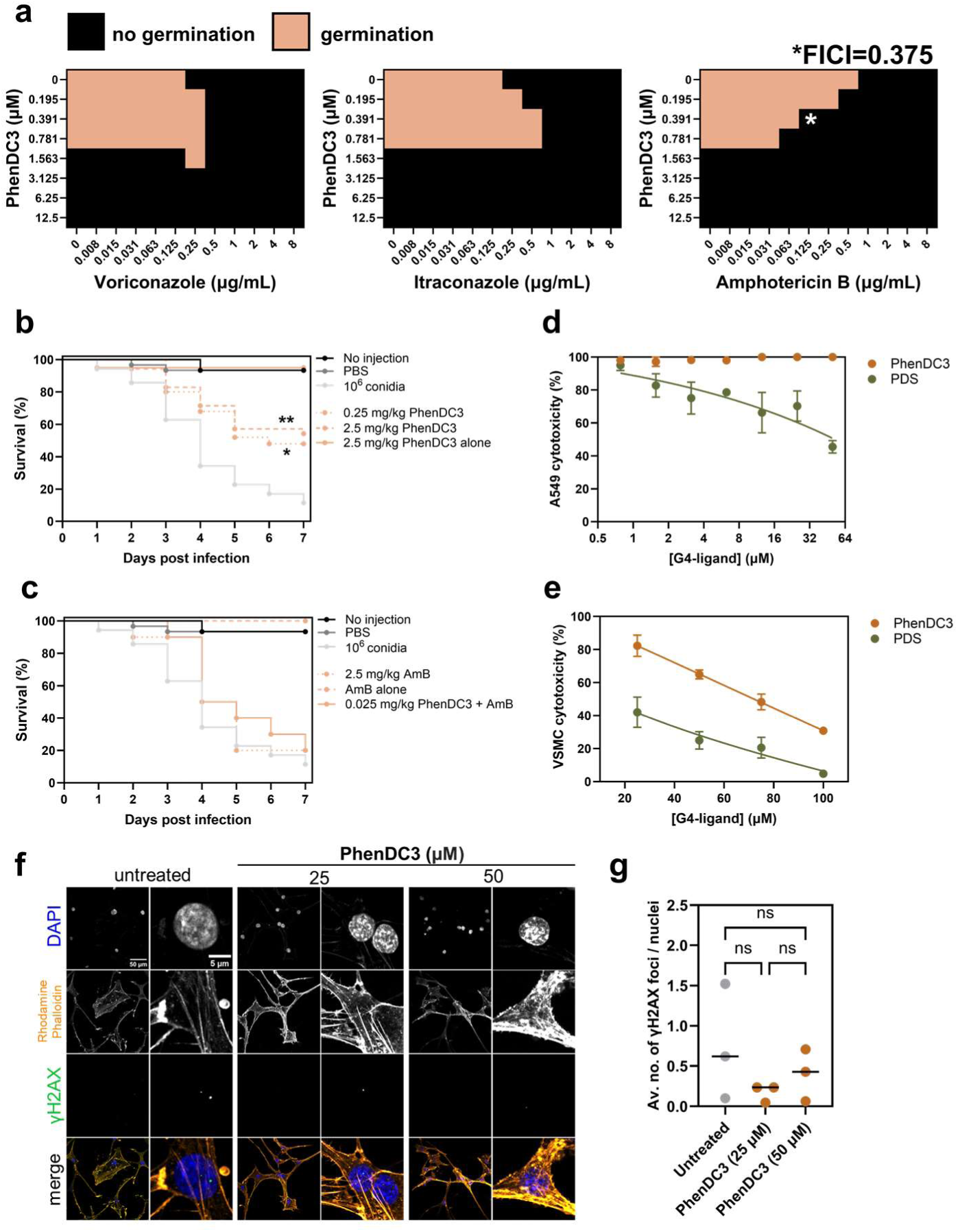
PhenDC3 synergises with amphotericin B, is protective in a *G. mellonella* infection model, and is well tolerated by human cells. (**a**) *A. fumigatus* conidia were treated with PhenDC3 in combination with the antifungals amphotericin B, itraconazole, or voriconazole at the concentrations indicated for 24 h in a checkerboard assay. The black cells indicate where germination was completely prevented. A fractional inhibitory concentration index (FICI) below 0.5 indicates synergy. (**b and c**) The percentage survival of *G. mellonella* larvae which were injected with PBS, 2.5 mg/kg amphotericin B (AmB) alone, 2.5 mg/kg PhenDC3 alone, or 1×10^6^ *A. fumigatus* conidia alone was monitored for 7-days. In some groups, larvae infected with 1×10^6^ *A. fumigatus* conidia were treated with 0.25 mg/kg PhenDC3, 2.5 mg/kg PhenDC3, 2.5 mg/kg AmB, or a combination of PhenDC3 and AmB (both at 0.025 mg/kg) and survival was monitored for 7-days. Data are representative of three biological replicates with 10 larvae per treatment group (30 total). Data for both figures are from the same experiments, but the figure has been split into two for clarity. Statistical significance was determined by Kaplan-Meier survival analyses (*p*=0.0018 (**) and *p*=0.0254 (*) for 0.25 mg/kg and 2.5 mg/kg PhenDC3 versus conidial infection alone). The viability of (**d**) the human A549 lung adenocarcinoma cell line or (**e**) primary human vascular smooth muscle cells was quantified following treatment with a concentration range of PhenDC3 or PDS for 24 h. Metabolism was measured using resazurin and normalised to the untreated control. (**f**) Representative confocal microscopy images of primary human vascular smooth muscle cells treated with 25 µM or 50 µM PhenDC3 for 24 h to quantify γH2AX foci, which are indicative of DNA damage. (**g**) The average number of γH2AX foci per nucleus following treatment with 25 µM or 50 µM for 24 h. Images were obtained using a Zeiss LSM 980 laser scanning confocal microscope equipped with an Airyscan 2 detector using the 20x objective. γH2AX foci were quantified using Fiji 1.53t. Experiments were performed in biological triplicate. Each data point represents the mean of three technical replicates. Statistical significance was determined by a Brown-Forsythe and Welch ANOVA test with Dunnett’s T3 multiple comparisons test (ns: not significant).

*Galleria mellonella* larvae (Greater wax-moth) have an innate immune system remarkably like humans and represent a useful model to study fungal infections *in vivo.*^25^ Uninfected *G. mellonella* larvae which were not injected, injected with PBS alone, or injected with 2.5 mg/kg PhenDC3 (equivalent to 4.91 µM) or 2.5 mg/kg amphotericin B (equivalent to 4.51 µM) displayed ≥90% viability over the 7-day experimentation period (**Fig. 4b, c)**. Larvae inoculated with 1×10^6^ *A. fumigatus* conidia alone displayed 89.6% mortality after 7 days. In larvae infected with *A. fumigatus,* 2.5 mg/kg amphotericin B, delayed larval death (50% versus 34% survival at day 4 for amphotericin B and control, respectively) and caused a small decrease in overall mortality at day 7 (9.6%; *p*>0.05). Notably, 0.25 mg/kg PhenDC3 (equivalent to 491 nM) also delayed larval death (68% versus 34% survival at day 4 for PhenDC3 and control, respectively) but reduced mortality at day 7 by 52% (*p*=0.0018). This concentration was more effective than 2.5 mg/kg PhenDC3 (36% survival at day 7; *p*=0.0254). As we found that PhenDC3 and amphotericin B could synergise *in vitro,* we tested whether this combination retained efficacy *in vivo.* For these experiments, we combined 10-fold less PhenDC3 with 100-fold less amphotericin B (both at a concentration of 0.025 mg/kg; equivalent to 49.1 nM and 45.1 nM, respectively). We found that this combination was equally as efficacious as amphotericin B alone at 2.5 mg/kg in this study (20% survival after 7 days) but was more effective at delaying larval death (68% versus 50% survival at day 4 for PhenDC3 and amphotericin B combined, versus amphotericin B alone).

PhenDC3 was highly effective against *A. fumigatus*, but G4s can be found in all organisms, including humans, so we next assessed the cytotoxic effects of G4-ligands against human cells. We assessed the cytotoxicity of PhenDC3 and PDS against the A549 type II lung alveolar epithelial cell line (representative of cells in the lung) and primary human vascular smooth muscle cells (representative of the predominant cell type in arteries and veins). Both PhenDC3 and PDS were well tolerated by A549 cells. There was no cytotoxicity observed in A549 cells for PhenDC3 up to 50 μM and the ED_50_ for PDS was ∼ 50 μM (**Fig. 4d)**. Conversely, PhenDC3 had much lower toxicity against smooth muscle cells comparative to PDS (ED_50_ of 72.8 μM for PhenDC3 versus 13.1 μM for PDS) (**Fig. 4e**).

Stabilising G4s has the potential to cause DNA damage and genome instability.^26^ We assessed the amount of DNA damage caused by PhenDC3 in the primary vascular smooth muscle cells by quantifying the number of γH2AX foci, indicative of double-strand breaks.^27^ We found that there was no significant difference in the number of γH2AX foci between control cells (mean 0.75 foci ± 0.42 SEM per nuclei) and cells treated with 25 μM (mean 0.17 foci ± 0.06 SEM per nuclei) or 50 μM (mean 0.40 foci ± 0.19 SEM per nuclei) PhenDC3 (**Fig. 4f, g**). PDS disrupted cell structure and caused cells to rupture, and DNA damage could not be quantified accurately in these cells (**Supplementary** Fig. 8).

### Prolonged G4 stabilisation can cause cross-resistance to current antifungals

Finally, we explored the development of resistance to PhenDC3. First, we attempted to determine how quickly resistance to itraconazole developed in our clinical isolate. *A. fumigatus* conidia were exposed to an MIC_50_ concentration of itraconazole for 48 h before conidia were sub-cultured on solid agar for 5 days and treated again with itraconazole for 48 h (**Fig. 5a**). This process was repeated until resistance emerged. We found that it took 5 passages (∼35 days experimental time) to observe itraconazole resistance (MIC_90_ > 32 µg/mL; **Fig. 5b**). We also found that these itraconazole-resistant isolates displayed some cross resistance to voriconazole (MIC_90_ = 16 µg/mL; **Fig. 5c**) but the MIC_90_ values for posaconazole were comparable between WT and the itraconazole-resistant isolate (MIC_90_ = 1 µg/mL; **Fig. 5d**).

**Figure 5.**
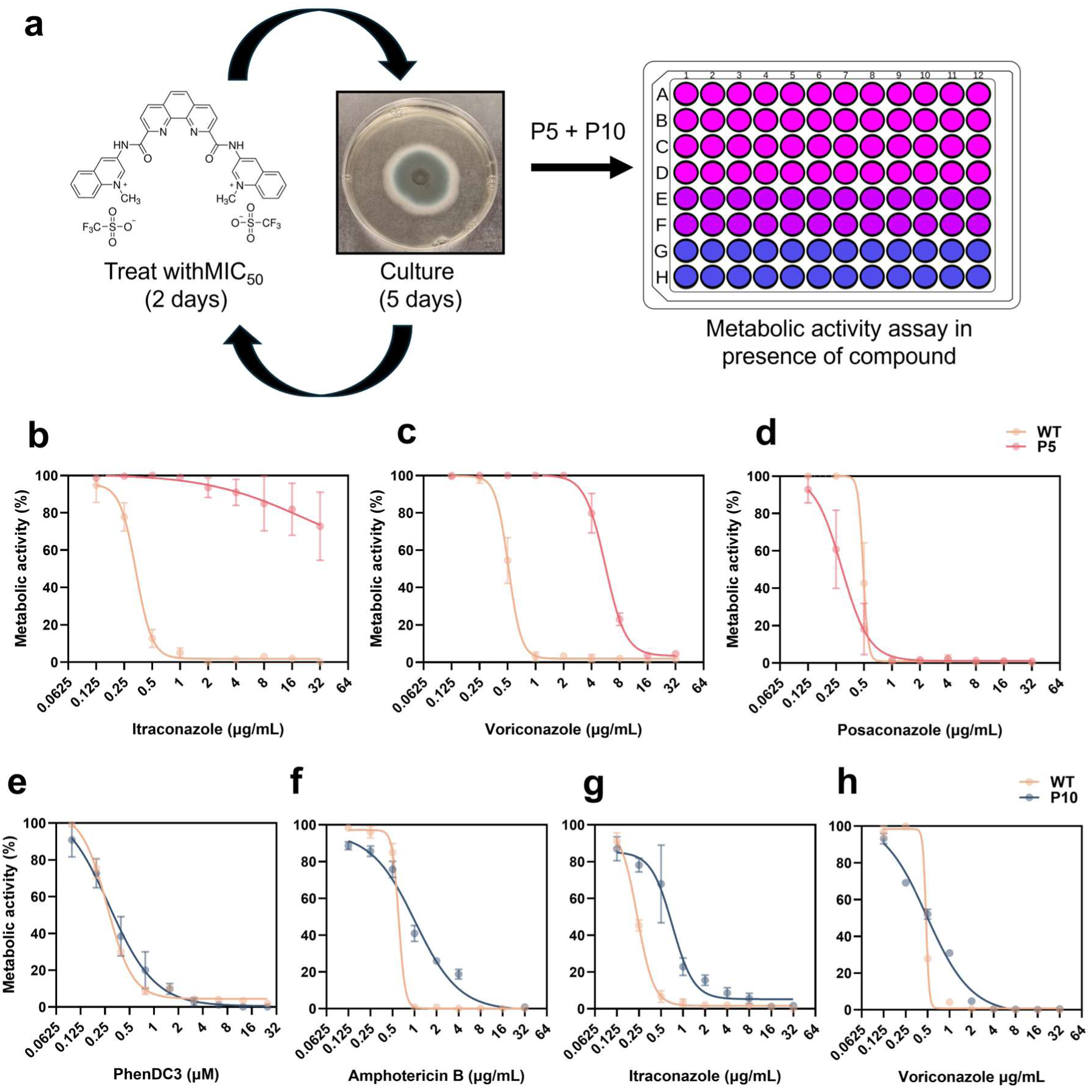
PhenDC3 resistance does not emerge within 70 days of treatment but does induce cross-resistance to current antifungals. (**a**) *A. fumigatus* conidia were treated with an MIC50 concentration of itraconazole for 2 days prior to subculture on SDA for 5 days. This was repeated 5 times (p5; 35 days). This itraconazole-treated isolate (p5) was treated with (**b**) itraconazole, (**c**) voriconazole or (**d**) posaconazole for 48 h to determine azole resistance. Concomitantly, *A. fumigatus* conidia were treated with an MIC50 concentration of PhenDC3 for 2 days prior to subculture on SDA for 5 days. This was repeated 10 times (p10; 70 days). This PhenDC3-treated isolate (p10) was treated with (**e**) PhenDC3, (**f**) amphotericin B, (**g**) itraconazole, or (**h**) voriconazole for 48 h to determine cross-resistance. Metabolic activity, quantified using resazurin, was calculated as percentage of the untreated control. Experiments are in biological triplicate with each data point consisting of the mean of three technical replicates ± SEM.

Conversely, we performed the same experiments with PhenDC3 and found no significant resistance had developed after 10 passages (∼70 days) (**Fig. 5e**). However, within this timeframe, some cross-resistance to amphotericin B, itraconazole, and voriconazole had emerged (**Fig. 5f-h**).

## Discussion

This work provided a comprehensive comparison between the impact of the G4-ligands PhenDC3 and PDS on fungal biology. We found that these ligands both exerted antifungal activities but in very different ways. Further, we showed that G4-ligands had the potential to be used therapeutically but additional in-depth studies are essential to fully realise the potential of G4s as a drug target in fungi. Additionally, we demonstrated the structural flexibility of fungal DNA to form a wide range of secondary structures that are often overlooked when considering the effects of drugs on the biological responses of fungi. This included two unusual G4s in *cyp51A* which were stabilised by acidic pH. Taken together, we highlight both the potential of G4s as novel drug targets and allude to the biological importance of non-canonical nucleic acids structures within fungal biology.

Although both PhenDC3 and PDS are designed to bind to G4s, their antifungal mechanism of action was very different. The way in which these ligands interact with DNA might provide some insight. We found that PhenDC3 caused the refolding of the A1-G and A2-G sequences into antiparallel G4 structures. Indeed, PhenDC3 has previously been shown to induce refolding of a hybrid G4 in the telomeres to an antiparallel chair-type structure by intercalating into the DNA.^28^ This is important to note as parallel G4s act as a substrate for telomerase binding, but antiparallel or hybrid G4s are poorly utilised as substrates for telomerase.^29–32^ It is possible that PhenDC3’s capability of inducing the non-native antiparallel form of the G4 prevents protein binding, and inhibits essential pathways in fungi. There were significant differences between the antifungal potencies of the G4-ligands between different species. This is suggestive that G4-ligands may be preventing the function of a specific gene (or set of genes) critical for the survival of these fungi. Indeed, through our analyses, we have found G4s to be present in some essential genes within *A. fumigatus,* but we need to explore this further.

Alternatively, accessibility to the G4s may differ between species as chromatin structure has a strong influence on G4 formation. Most endogenous G4s can be found mapped to nucleosome-depleted, open regions of chromatin, and certain G4s may be inaccessible to G4-ligands across species.^33^ Although this doesn’t explain the faster and wider activity of PDS. It may be that PDS exerts its antifungal activity non-specifically and could simply be due to the promotion of genome instability and inhibition of DNA replication as observed in the fellow yeast *Saccharomyces cerevisiae.*^34–36^

Unusually, we also identified two G4s in *cyp51A* and *cyp51B* which transitioned to stable mixed species G4s at pH 5.5. It is widely appreciated that the cytosine-rich iM is readily stabilised by acidic pH, but acidity does not typically affect G4s.^13,37^ However, there has been at least one previously reported case of an acid-stabilised G4 that can form within the telomeric region, but reports are rare.^38^ We are currently unsure why this occurs in these *cyp* genes and what implications this might have on the function of both Cyp51A and Cyp51B, but both proteins are implicated in azole resistance and acidity has been shown to affect azole resistance in fungi.^39,40^ Therefore, the formation of the mixed G4 in acidic conditions might be a contributing factor to azole resistance and should be explored further.

Notably, global stabilisation of G4s for a prolonged duration (∼70 days) induced cross resistance to current antifungals. As aforementioned, G4s can contribute to genome instability, and this can induce DNA damage leading to mutations.^34–36^ Our previous bioinformatic study identified G4-forming sequences in several genes implicated in antifungal resistance, including *cyp51A, cyp51B, atrR, abcC,* and *hmg1.*^21^ Therefore, prolonged global stabilisation could lead to the increased stability of these G4s, causing mutations, and promoting resistance. This highlights the necessity of studying these non-canonical structures and their roles in fungal biology further, particularly if G4s can contribute to the acquisition of resistance.

These findings offer the first insights into the potential therapeutic applications of stabilising distinct secondary structures within the genomes of fungal pathogens, with further implications for genomic stability, drug resistance, and regulation in *Aspergillus* and *Candida* species. However, further study is required to identify the specific targets of G4-stabilisers and to elucidate the biological roles of G4s in fungi to help develop novel antifungals targeting specific G4s.

## Online Methods

### Fungal isolates and culture

*Aspergillus* spp. and *Candida* spp. (**Supplementary Table 1**) were routinely grown on Sabouraud Dextrose Agar (SDA; Oxoid, UK) for 3-5 days for *Aspergillus* spp. and 1-2 days for *Candida* spp. at 37 °C in T-75 tissue culture flasks (Thermo Fisher, UK). *Aspergillus* conidia were harvested using PBS-Tween 20 (0.05%) and large hyphal fragments were removed by passing the harvested fungi through Miracloth (VWR Scientific, USA). Conidia were pelleted at 3000*g* for 5 min and washed twice in sterile PBS. *Candida* was harvested by inoculating 3-5 distinct colonies in sterile water.

### Tissue culture

The type II alveolar epithelial/adenocarcinoma cell line, A549 (a kind gift from Dr Anastasia Sobolewski, University of East Anglia) was maintained in DMEM F-12 supplemented with 2 mM L-glutamine (Gibco, Fisher Scientific, UK), 10% foetal bovine serum (Fisher Scientific, UK), and 100 U/mL penicillin plus 100 µg/mL streptomycin (Fisher Scientific, UK). Primary vascular smooth muscle cells were purchased from Cell Applications Inc. (USA; Cat#354-05a). Standard culture (passage 3-9) was performed as previously described.^41,42^ Cells were maintained at 37 °C in a humidified incubator supplied with 5% CO_2._

### Antifungal susceptibility testing

Antifungal susceptibility assays were conducted according to adapted Clinical and Laboratory Standards Institute (CLSI) reference methods: M27Ed4 for *Candida* spp. and M38Ed3 for *Aspergillus* spp. Conidia or yeast were incubated with DMSO as a vehicle control, amphotericin B (2 µg/mL; Merck, UK) as a positive control, or the G4-ligands PhenDC3 and PDS (both purchased from Merck, UK) in RPMI-1640 growth medium (Gibco, UK) buffered with 0.165 M

MOPS (3-(*N-*morpholino)propanesulfonic acid) buffer (pH 7.0) at 37 °C for 24 h or 48 h where indicated. The final concentration of DMSO in each instance was <0.05%. Following incubation, 10 μg/mL resazurin (Merck, UK) in PBS was added to each well and incubated for 4 h for *Candida* spp. or 24 h for *Aspergillus* spp. at 37 °C. The ability of metabolically active cells to convert resazurin to the fluorescent product resorufin was quantified with a CLARIOstar plate reader (BMG Labtech, UK) using an excitation wavelength of 530 nm and an emission wavelength of 590 nm.

For checkerboard assays, *Aspergillus* spp. conidia (1×10^6^) were added to each well of a 96-well plate containing RPMI-1640 growth medium buffered with 0.165 M MOPS (pH 7.0). PhenDC3 (0.09 to 6.25 µM), amphotericin B, itraconazole, or voriconazole (all at concentrations from 0.031 to 8 µg/mL and purchased from Merck, UK) were added to conidia for 24 h alone or in combination at 37 °C. Metabolic activity was determined as above following incubation with 10 μg/mL resazurin for 24 h. Concentrations exhibiting a >50% decrease in viability was used as a cut-off for activity in the checkerboard assays and the fractional inhibitory concentration index was determined using the following equation:

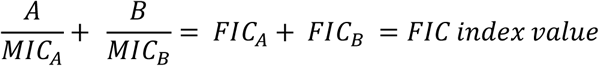

For experiments exploring resistance/tolerance, *A. fumigatus* was sub-cultured on SDA for 5 days as above, prior to treatment with an MIC_50_ concentration of PhenDC3 or itraconazole for 48 h in RPMI-1640 media containing 0.165 M MOPS buffer (pH 7.0) at 37 °C. This process was repeated for 70 days.

In all instances, the background fluorescence after amphotericin B treatment was subtracted from the test samples, as this value represented the intrinsic fluorescence of the dye in the absence of viable fungi (0% viability). Metabolic activity was calculated as a percentage of the untreated control (100% viability).

### Scanning electron microscopy (SEM)

Conidia (1×10^6^) were treated with 12.5 µM PhenDC3 for 4 h at 37 °C. Following incubation, conidia were pelleted in PBS containing 0.05% Tween-20 at 3000*g* for 5 min and fixed in 2.5% glutaraldehyde for 3 h. Fixed conidia were washed and resuspended in 0.05 M sodium cacodylate buffer (pH 7.3), transferred onto 13 mm poly-L-lysine coated coverslips (0.01% w/v), and left to settle for 20 min. Conidia were post-fixed in 1% (v/v) OsO_4_ for 1 h at room temperature. The OsO_4_ fixation was followed by three 15 min washes in distilled water before beginning the ethanol dehydration series (25%, 50%, 75%, 95% and two changes of 100% ethanol, each for an hour) and left to air dry overnight. The sample was then plunged into a liquid nitrogen slush and transferred under vacuum onto a PP3010T Cryo-SEM preparation system (Quorum Technologies, UK) at −140°C. The sample was sublimated for 2 min at −90 °C before cooling again to −140 °C and sputter-coating with platinum for 1 min at 10 mA. The sample was introduced onto the cryo-stage in a Gemini 300 scanning electron microscope (Carl Zeiss, Germany) and viewed at 2 kV.

### Galleria mellonella infection model

Larvae of *Galleria mellonella* (Live Foods Direct, UK) were used for all experiments. Larvae were maintained in their box in the dark at room temperature prior to use. Only larvae weighing between 100 and 300 mg, with normal mobility, and uniform colour were selected for the experiments. Larvae were randomly distributed into groups of 10. Larvae were administered 1×10^6^ *A. fumigatus* conidia in sterile PBS via injection in the ventral side of the last proleg using a Hamilton® syringe. Groups of larvae injected with PBS alone, 2.5 mg/kg PhenDC3 alone, 2.5 mg/kg amphotericin B alone, or not injected at all were included as controls. Two hours post-infection, either 2.5 mg/kg amphotericin B, 0.25 mg/kg PhenDC3, or a combination of 0.25 mg/kg PhenDC3 and 0.25 mg/kg amphotericin B were administered via injection into the haemocoel. Infected larvae were stored in the dark at 37 °C and mortality was monitored for 7 days. Larvae were considered dead when they melanised and stopped responding to stimulation. All treatments were prepared in sterile PBS and administered in a 10 µL inoculum. Experiments were performed in triplicate and results pooled.

### Cytotoxicity assays

To assess the cytotoxicity of PhenDC3 and PDS against A549 cells and primary vascular smooth muscle cells, cells were plated at a density of 2 x 10^4^ cells/well in their respective growth buffers in a 96-well plate and incubated at 37 °C and 5% CO_2_ for 24 h. The following day, the growth media was aspirated and replaced with growth media containing PhenDC3 or PDS at a concentration range from 0.7-100 µM. Cells were incubated for a further 24 h prior to the addition of resazurin (10 μg/mL) and incubation at 37 °C and 5% CO_2_ for 2 h. The ability of metabolically active cells to convert resazurin to the fluorescent product resorufin was quantified with a CLARIOstar plate reader using an excitation wavelength of 530 nm and an emission wavelength of 590 nm.

### QUMA-1 assay

Fungi were treated with 12.5 µM of the G4-ligands PhenDC3 or PDS in a glass bottomed µ-Plate 96-well black plate (Ibidi, Germany) for 18 h prior to labelling with QUMA-1 (Merck, UK). Following treatment, fungi were washed in PBS + 0.05% Tween-20 and resuspended in PBS containing 1 µM Hoechst 33342 (Merck, UK) and 1 µM QUMA-1. Live fungi were stained at 37 °C for 1 h and imaged immediately on a Zeiss LSM 980 laser scanning confocal microscope equipped with an Airyscan 2 detector (Carl Zeiss, Germany) using a 100x oil objective. The average foci per 10 µm^2^ in all fungi from five random images per treatment from three independent experiments were quantified using Fiji 1.53t.

### RT-qPCR

Total RNA was extracted from cells using TRIzol reagent (Invitrogen, UK) with chloroform separation and isopropanol precipitation. RNA was quantified using Luna^®^ Universal One-Step RT-qPCR Kit (New England Biolabs, UK). Briefly, 1 μg of isolated total RNA, oligonucleotides (Cyp51a_F, Cyp51a_R, TubA_F, TubA_R), and Luna^®^ Reaction Mix were added to a final volume of 20 μl. Reverse transcription was carried out at 55 °C for 10 minutes, followed by the initial denaturation at 95°C for 1 minute. Then, 40 thermocycles at 95 °C for 1 minutes and 60 °C for 30 seconds were performed. PCR quantification was conducted using the 2−ΔΔCT method and normalized to TubA expression.

The primers used were as follows:

Cyp51a_F - TGCGTGCAGAGAAAAGTATGG

Cyp51a_R - AGACCTCTTCCGCATTGACA

TubA_F - CTCCGTTCCTGAGTTGACCC

TubA_R - CACGGAAAATGGCAGAGCAG

### DNA damage

Vascular smooth muscle cells were seeded in 6-well plates in VSMC growth media for 24 h at 37 °C and 5% CO_2_. Following incubation, the growth media was aspirated and replaced with growth media containing 25 μM or 50 μM PhenDC3/PDS and incubated for 24 h at 37 °C and 5% CO_2_. The media was removed, and cells were washed with PBS prior to fixation with paraformaldehyde for 10 min and permeabilization with 0.5% NP-40 (Thermo Fisher, UK) for 5 min. Cells were blocked with 3% bovine serum albumin (Merck, UK) for 1 h and stained with a γH2AX primary antibody (1:100; Cell Signalling Technology, UK) overnight at 4 °C. The primary antibody was washed away with PBS prior to the addition of the Alexa Fluor anti-mouse 488 secondary antibody (1:400; Invitrogen, UK) and rhodamine phalloidin stain (1:400; Invitrogen, UK). Cells were incubated in the dark at room temperature for 2 h prior to washing three times with PBS and mounting on microscope slides using DAPI mounting media (Vector Laboratories, UK). Images were obtained using a Zeiss LSM 980 laser scanning confocal microscope equipped with an Airyscan 2 detector (Carl Zeiss, Germany) using the 20x objective. Images were analysed using Fiji 1.53t.

### Oligonucleotides

All DNA sequences used in this study were synthesised and reverse-phase high-performance liquid chromatography (HPLC)-purified by Eurogentec (Eurogentec, Belgium). The oligonucleotides were dissolved in ultra-pure water to a final concentration of 1 mM, and their concentrations were confirmed using a NanoDrop (Thermo Fisher, UK). Sodium cacodylate was used in line with previous studies at 10 mM with 100 mM of either LiCl, NaCl, or KCl at the specified pH.^43^ For biophysical analysis, samples were annealed by heating at 95 °C for 5 min in a heating block, followed by cooling to room temperature overnight.

### Circular Dichroism (CD) Spectroscopy

The CD spectra of selected fungal *cyp51* sequences at various pH values were recorded using a JASCO J-1500 spectropolarimeter under a constant flow of nitrogen gas (JASCO, UK). Samples were diluted to 10 μM in 10 mM sodium cacodylate buffer containing 100 mM LiCl or 100 mM NaCl at pH 7, and 100 mM KCl buffer at pH values ranging from 4 to 8. Four scans were accumulated for each sample and corresponding buffer blank over a wavelength range of 230–320 nm at 25 °C. The instrument settings were as follows: data pitch of 0.5 nm, scanning speed of 200 nm/min, response time of 1 second, bandwidth of 1 nm, and sensitivity of 200 mdeg. Data were zero-corrected at 320 nm. The transitional pH (pH_T_) was determined from at least two independent experiments by identifying the inflection point of the sigmoidal fit of ellipticity measured at 288 nm or 295 nm, as appropriate for the sequence and pH range.

CD spectroscopy was also used to measure any ligand-induced effects on the fungal *cyp51* sequences (A1-G, A2-G and B-G). All samples were diluted to 10 μM in 10 mM sodium cacodylate buffer containing 100 mM KCl at pH 7.0. Four scans were accumulated over the wavelength range of 230–320 nm at 25°C with the same instrument settings as above. Ligands PhenDC3 and pyridostatin were titrated stepwise into the DNA samples via consecutive additions, with final ligand concentration ranges of 0–90 μM for PhenDC3 and 0– 170 μM for PDS. Titrations were performed in duplicate, and data were corrected for solvent effects, smoothed using the Savitzky–Golay method with a 10-point window, and zero-corrected at 320 nm. Data analysis was conducted using OriginLab software (version 2023B; OriginLab Corporation, USA). The sigmoidal curves of ellipticity at 295 nm versus ligand concentration were fitted to the Hill equation to obtain the Hill coefficient (n) and K, representing the ligand concentration required to achieve 50% reduction of molar ellipticity.

### UV Melting/Annealing and Thermal Difference Spectroscopy (TDS)

UV melting/annealing and TDS experiments were performed using a Jasco V-750 UV–Vis spectrophotometer (JASCO, UK). Samples were prepared at a concentration of 5 μM in 10 mM sodium cacodylate buffer containing 100 mM LiCl, 100 mM NaCl, or 100 mM KCl at pH 7, and 100 mM KCl at pH 5.5. For the UV melting and annealing experiments, absorbance was recorded at 260 nm and 295 nm every 1 °C over three thermal cycles. Samples were initially held at 4 °C for 10 min, followed by a gradual increase to 95 °C at a rate of 0.5 °C/min (melting phase). Upon reaching 95 °C, samples were held for 10 min before reversing the process to return to 4 °C (annealing phase). At each temperature interval, samples were equilibrated for 5 min before measurement. The most prominent melting temperature (T_m_) and annealing temperature (T_A_) were determined using the first derivative method for each cycle.

Thermal difference spectra were obtained by measuring the absorbance spectra from 230– 320 nm after equilibrating the samples at 4 °C for 10 min (folded DNA) and at 95 °C for 10 min (unfolded DNA). The TDS signatures were calculated by subtracting the absorbance spectra of the folded structures from those of the unfolded structures, zero-corrected at 320 nm, and normalised to the maximum absorbance.

## Statistics

The sample sizes (*n*), the probability (*p*) value and statistical test used to analyse each experiment can be found in the figure legends. Statistical significance was determined by paired Student’s *t*-test to compare two paired groups. To compare multiple groups a Kruskal-Wallis with Dunn’s multiple comparisons or Brown-Forsythe and Welch ANOVA with Dunnett’s T3 multiple comparisons was conducted. Survival analyses were analysed using Kaplan-Meier survival plots. Data was prepared using GraphPad Prism (version 10.0.3) and OriginPro (version 2023B). Data from most experiments are presented as mean values ± SEM. Data of UV spectroscopy are shown as mean ± SD. *p<*0.05 was considered statistically significant. Data is representative of at least two independent experiments.

## Acknowledgements

We are thankful for the support of A. Sobolewski (University of East Anglia, Norwich) and the A549 cells used in this study. This work was supported by an Academy of Medical Sciences Springboard Award (SBF008\1046 to S.B. and G.M.), BBSRC New Investigator Award (BB/Y005058/1 to S.B. and I.S.), a Thomas Marns Scholarship (to F.M.), a BBSRC grant (BB/W001616/1 to Z.A.E.W.), and a Wellcome Trust grant (226408/Z/22/Z to N.v.R.).

## Author information

These authors contributed equally: Georgie Middleton, Fuad Muhamud, and Isabelle Storer. Norman van Rhijn has competing interests and has received speaker honoraria from Napp/Mundipharma.

## Author contributions

S.B., Z.A.E.W., D.W. conceived and designed the study. S.B., G.M. and I.S. prepared the initial draft of the manuscript. All authors have contributed to the writing and editing of the final manuscript. G.M., F.M., I.S., performed most of the experiments and data analysis. A.W-G., F.W., E.B., A.B., O.S., performed some of the experiments and data analysis. A.A., S.S., J.M., B.L., N.v.R., A.d.S.D., T.F., and D.W. provided critical materials, experimental guidance, or essential technical support for the success of the study.

## References

1 Denning, D. W. Global incidence and mortality of severe fungal disease. Lancet Infect Dis 24, e428–e438 (2024). 10.1016/s1473-3099(23)00692-8

2 Case, N. T. et al. The future of fungi: threats and opportunities. G3 (Bethesda) 12 (2022). 10.1093/g3journal/jkac224

3 Seagle, E. E., Williams, S. L. & Chiller, T. M. Recent Trends in the Epidemiology of Fungal Infections. Infect Dis Clin North Am 35, 237–260 (2021). 10.1016/j.idc.2021.03.001

4 Fisher, M. C. & Denning, D. W. The WHO fungal priority pathogens list as a game-changer. Nat Rev Microbiol 21, 211–212 (2023). 10.1038/s41579-023-00861-x

5 Snelders, E. et al. Triazole fungicides can induce cross-resistance to medical triazoles in Aspergillus fumigatus. PLoS One 7, e31801 (2012). 10.1371/journal.pone.0031801

6 Fraaije, B., Atkins, S., Hanley, S., Macdonald, A. & Lucas, J. The Multi-Fungicide Resistance Status of Aspergillus fumigatus Populations in Arable Soils and the Wider European Environment. Front Microbiol 11, 599233 (2020). 10.3389/fmicb.2020.599233

7 Rhodes, J. et al. Population genomics confirms acquisition of drug-resistant Aspergillus fumigatus infection by humans from the environment. Nat Microbiol 7, 663–674 (2022). 10.1038/s41564-022-01091-2

8 Bromley, M. J. et al. Occurrence of azole-resistant species of Aspergillus in the UK environment. J Glob Antimicrob Resist 2, 276–279 (2014). 10.1016/j.jgar.2014.05.004

9 van Rhijn, N. et al. Aspergillus fumigatus strains that evolve resistance to the agrochemical fungicide ipflufenoquin in vitro are also resistant to olorofim. Nat Microbiol 9, 29–34 (2024). 10.1038/s41564-023-01542-4

10 Varshney, D., Spiegel, J., Zyner, K., Tannahill, D. & Balasubramanian, S. The regulation and functions of DNA and RNA G-quadruplexes. Nat Rev Mol Cell Biol 21, 459–474 (2020). 10.1038/s41580-020-0236-x

11 Mergny, J. L., De Cian, A., Ghelab, A., Saccà, B. & Lacroix, L. Kinetics of tetramolecular quadruplexes. Nucleic Acids Res 33, 81–94 (2005). 10.1093/nar/gki148

12 Gehring, K., Leroy, J. L. & Guéron, M. A tetrameric DNA structure with protonated cytosine.cytosine base pairs. Nature 363, 561–565 (1993). 10.1038/363561a0

13 Day, H. A., Pavlou, P. & Waller, Z. A. i-Motif DNA: structure, stability and targeting with ligands. Bioorg Med Chem 22, 4407–4418 (2014). 10.1016/j.bmc.2014.05.047

14 Georgakopoulos-Soares, I. et al. Alternative splicing modulation by G-quadruplexes. Nat Commun 13, 2404 (2022). 10.1038/s41467-022-30071-7

15 Li, P. T. et al. Expression of the human telomerase reverse transcriptase gene is modulated by quadruplex formation in its first exon due to DNA methylation. J Biol Chem 292, 20859–20870 (2017). 10.1074/jbc.M117.808022

16 De Cian, A., Delemos, E., Mergny, J. L., Teulade-Fichou, M. P. & Monchaud, D. Highly efficient G-quadruplex recognition by bisquinolinium compounds. J Am Chem Soc 129, 1856–1857 (2007). 10.1021/ja067352b

17 Rodriguez, R. et al. A novel small molecule that alters shelterin integrity and triggers a DNA-damage response at telomeres. J Am Chem Soc 130, 15758–15759 (2008). 10.1021/ja805615w

18 Cebrián, R. et al. G-Quadruplex DNA as a Target in Pathogenic Bacteria: Efficacy of an Extended Naphthalene Diimide Ligand and Its Mode of Action. J Med Chem 65, 4752–4766 (2022). 10.1021/acs.jmedchem.1c01905

19 Ruggiero, E. & Richter, S. N. G-quadruplexes and G-quadruplex ligands: targets and tools in antiviral therapy. Nucleic Acids Res 46, 3270–3283 (2018). 10.1093/nar/gky187

20 Harris, L. M. et al. G-Quadruplex DNA Motifs in the Malaria Parasite Plasmodium falciparum and Their Potential as Novel Antimalarial Drug Targets. Antimicrob Agents Chemother 62 (2018). 10.1128/aac.01828-17

21 Warner, E. F., Bohálová, N., Brázda, V., Waller, Z. A. E. & Bidula, S. Analysis of putative quadruplex-forming sequences in fungal genomes: novel antifungal targets? Microb Genom 7 (2021). 10.1099/mgen.0.000570

22 Chen, X. C. et al. Tracking the Dynamic Folding and Unfolding of RNA G-Quadruplexes in Live Cells. Angew Chem Int Ed Engl 57, 4702–4706 (2018). 10.1002/anie.201801999

23 Mergny, J. L., Li, J., Lacroix, L., Amrane, S. & Chaires, J. B. Thermal difference spectra: a specific signature for nucleic acid structures. Nucleic Acids Res 33, e138 (2005). 10.1093/nar/gni134

24 Benabou, S., Mazzini, S., Aviñó, A., Eritja, R. & Gargallo, R. A pH-dependent bolt involving cytosine bases located in the lateral loops of antiparallel G-quadruplex structures within the SMARCA4 gene promotor. Sci Rep 9, 15807 (2019). 10.1038/s41598-019-52311-5

25 Giammarino, A., Bellucci, N. & Angiolella, L. Galleria mellonella as a Model for the Study of Fungal Pathogens: Advantages and Disadvantages. Pathogens 13 (2024). 10.3390/pathogens13030233

26 De Magis, A. et al. DNA damage and genome instability by G-quadruplex ligands are mediated by R loops in human cancer cells. Proc Natl Acad Sci U S A 116, 816–825 (2019). 10.1073/pnas.1810409116

27 Kuo, L. J. & Yang, L. X. Gamma-H2AX - a novel biomarker for DNA double-strand breaks. In Vivo 22, 305–309 (2008).

28 Ghosh, A., Trajkovski, M., Teulade-Fichou, M. P., Gabelica, V. & Plavec, J. Phen-DC(3) Induces Refolding of Human Telomeric DNA into a Chair-Type Antiparallel G-Quadruplex through Ligand Intercalation. Angew Chem Int Ed Engl 61, e202207384 (2022). 10.1002/anie.202207384

29 Zahler, A. M., Williamson, J. R., Cech, T. R. & Prescott, D. M. Inhibition of telomerase by G-quartet DNA structures. Nature 350, 718–720 (1991). 10.1038/350718a0

30 Zaug, A. J., Podell, E. R. & Cech, T. R. Human POT1 disrupts telomeric G-quadruplexes allowing telomerase extension in vitro. Proc Natl Acad Sci U S A 102, 10864–10869 (2005). 10.1073/pnas.0504744102

31 Paudel, B. P. et al. A mechanism for the extension and unfolding of parallel telomeric G-quadruplexes by human telomerase at single-molecule resolution. Elife 9 (2020). 10.7554/eLife.56428

32 El-Khoury, R. et al. Telomeric i-motifs and C-strands inhibit parallel G-quadruplex extension by telomerase. Nucleic Acids Res 51, 10395–10410 (2023). 10.1093/nar/gkad764

33 Shen, J. et al. Promoter G-quadruplex folding precedes transcription and is controlled by chromatin. Genome Biol 22, 143 (2021). 10.1186/s13059-021-02346-7

34 Piazza, A. et al. Genetic instability triggered by G-quadruplex interacting Phen-DC compounds in Saccharomyces cerevisiae. Nucleic Acids Res 38, 4337–4348 (2010). 10.1093/nar/gkq136

35 Lopes, J. et al. G-quadruplex-induced instability during leading-strand replication. Embo j 30, 4033–4046 (2011). 10.1038/emboj.2011.316

36 Paeschke, K. et al. Pif1 family helicases suppress genome instability at G-quadruplex motifs. Nature 497, 458–462 (2013). 10.1038/nature12149

37 König, S. L., Huppert, J. L., Sigel, R. K. & Evans, A. C. Distance-dependent duplex DNA destabilization proximal to G-quadruplex/i-motif sequences. Nucleic Acids Res 41, 7453–7461 (2013). 10.1093/nar/gkt476

38 Galer, P., Wang, B., Šket, P. & Plavec, J. Reversible pH Switch of Two-Quartet G-Quadruplexes Formed by Human Telomere. Angew Chem Int Ed Engl 55, 1993–1997 (2016). 10.1002/anie.201507569

39 Lourenço, A., Pedro, N. A., Salazar, S. B. & Mira, N. P. Effect of Acetic Acid and Lactic Acid at Low pH in Growth and Azole Resistance of Candida albicans and Candida glabrata. Front Microbiol 9, 3265 (2018). 10.3389/fmicb.2018.03265

40 Song, J. et al. Acidic/Alkaline Stress Mediates Responses to Azole Drugs and Oxidative Stress in Aspergillus fumigatus. Microbiol Spectr 10, e0199921 (2022). 10.1128/spectrum.01999-21

41 Ragnauth, C. D. et al. Prelamin A acts to accelerate smooth muscle cell senescence and is a novel biomarker of human vascular aging. Circulation 121, 2200–2210 (2010). 10.1161/circulationaha.109.902056

42 Warren, D. T. et al. Nesprin-2-dependent ERK1/2 compartmentalisation regulates the DNA damage response in vascular smooth muscle cell ageing. Cell Death Differ 22, 1540–1550 (2015). 10.1038/cdd.2015.12

43 Martella, M. et al. i-Motif formation and spontaneous deletions in human cells. Nucleic Acids Res 50, 3445–3455 (2022). 10.1093/nar/gkac158

