## Supplemental Figures for "G-quadruplexes represent promising new targets to overcome multidrug-resistant fungal infections"

**Supplementary Tables**

**Supplementary Table 1.** The fungal isolates used in this study and minimum inhibitory concentrations

| | | **$MIC_{50/90}$ (µM)** | **$MIC_{50/90}$ (µM)** |
|---|---|---|---|
| **Species** | **Strain** | **PhenDC3** | **PDS** |
| *A. fumigatus* | 22M7007854 | 0.83/1.56 | 6.96/12.50 |
| *A. fumigatus* | 22M7004177 | 0.40/0.78 | 7.93/12.50 |
| *A. fumigatus* | CEA10 | 0.89/3.13 | 5.38/6.25 |
| *A. fumigatus* | A1160+ | 1.13/3.13 | 3.31/6.25 |
| *A. fumigatus* | $TR_{34}$/L98H | 0.38/0.78 | 5.95/12.50 |
| *A. hiratsukae* | Clinical | 1.10/>12.50 | 3.98/6.25 |
| *A. udagawae* | Clinical | 0.69/6.25 | 6.09/12.50 |
| *A. flavus* | 22M8001325 | ND/ND | ND/ND |
| *A. niger* | ATCC16404 | ND/ND | 11.43/12.50 |
| *C. albicans* | SC5314 | ND/ND | 5.94/12.5 |
| *C. auris* | NCPF8985 | 30.27/50 | 5.48/6.25 |
| *C. glabrata* | CBS-128 | ND/ND | 3.94/6.25 |

ND = Not determined

**Supplementary Figures**

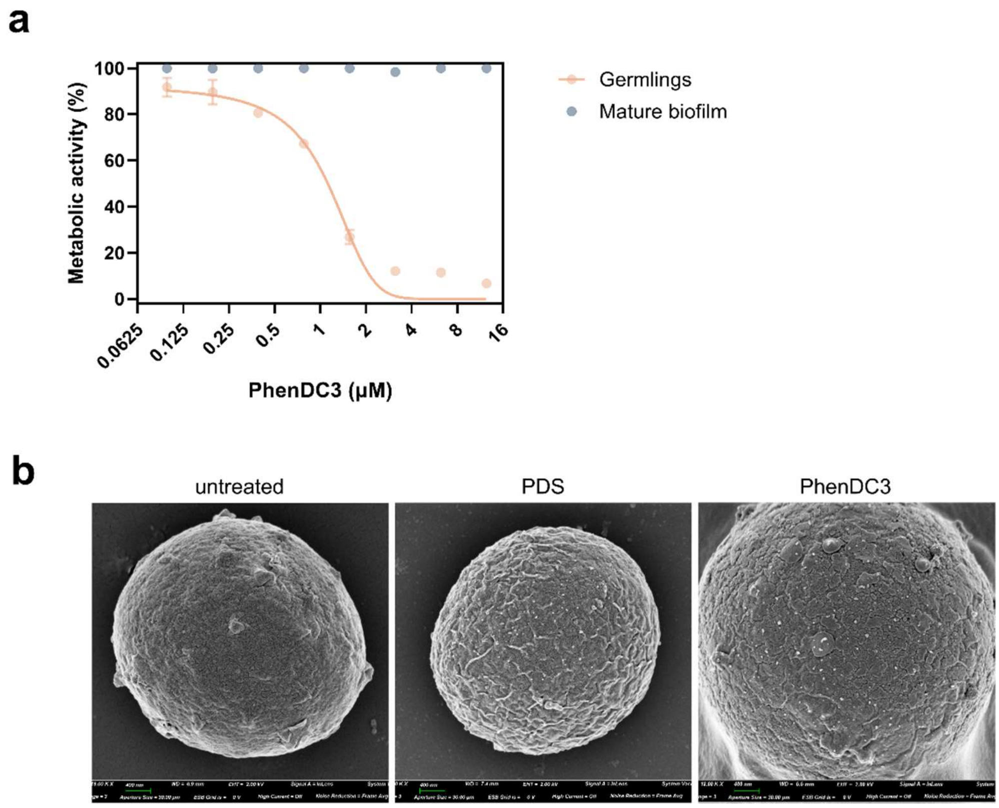


**Supplementary Fig. 1.** (**a**) *A. fumigatus* germlings (6 h of growth) or mature biofilms (<24 h growth) were treated with the indicated concentrations of PhenDC3 for 48 h at 37 °C. Metabolic activity was calculated as percentage of the untreated control. Experiments are in biological triplicate with each data point consisting of the mean of three technical replicates ± SEM. (**b**) *A. fumigatus* conidia were treated with 12.5 µM PDS or PhenDC3 for 24 h prior to scanning electron microscopy. The sample was sputter-coated with platinum for 1 min at 10 mA before imaging on a Zeiss Gemini 300 scanning electron microscope and viewed at 2 kV.

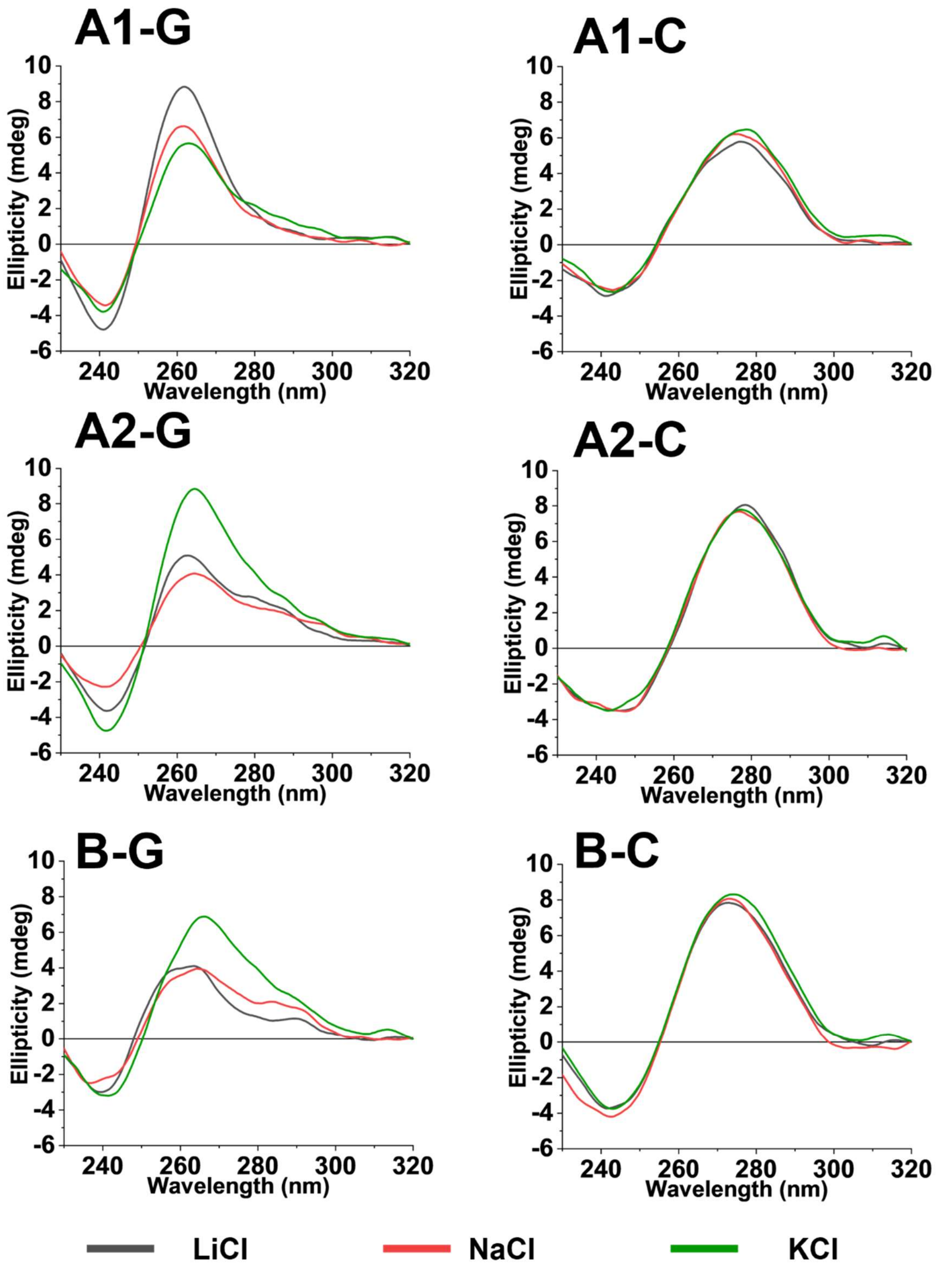


**Supplementary Fig. 2.** CD Spectroscopy of the *cyp51* sequences A1-G, A1-C, A2-G, A2-C, B-G, and B-C. DNA oligonucleotides (10 μM) were annealed in 10 mM sodium cacodylate buffer containing 100 mM LiCl, NaCl, or KCl (as indicated) at pH 7.0.

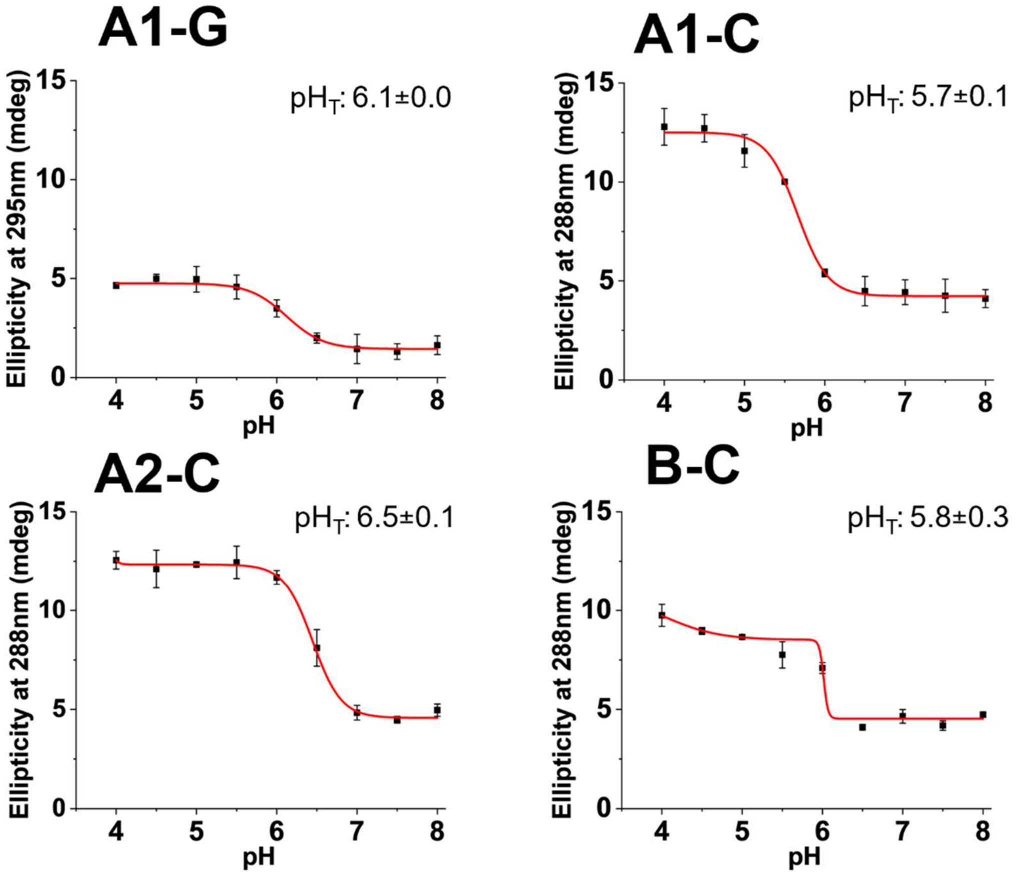


**Supplementary Fig. 3.** Corresponding plot ellipticity for the CD experiments in Fig. 1c. at 288nm at the different pH. This plot was used to determine the transitional pH ($pH_T$) from the inflection point of a Boltzmann sigmoidal.

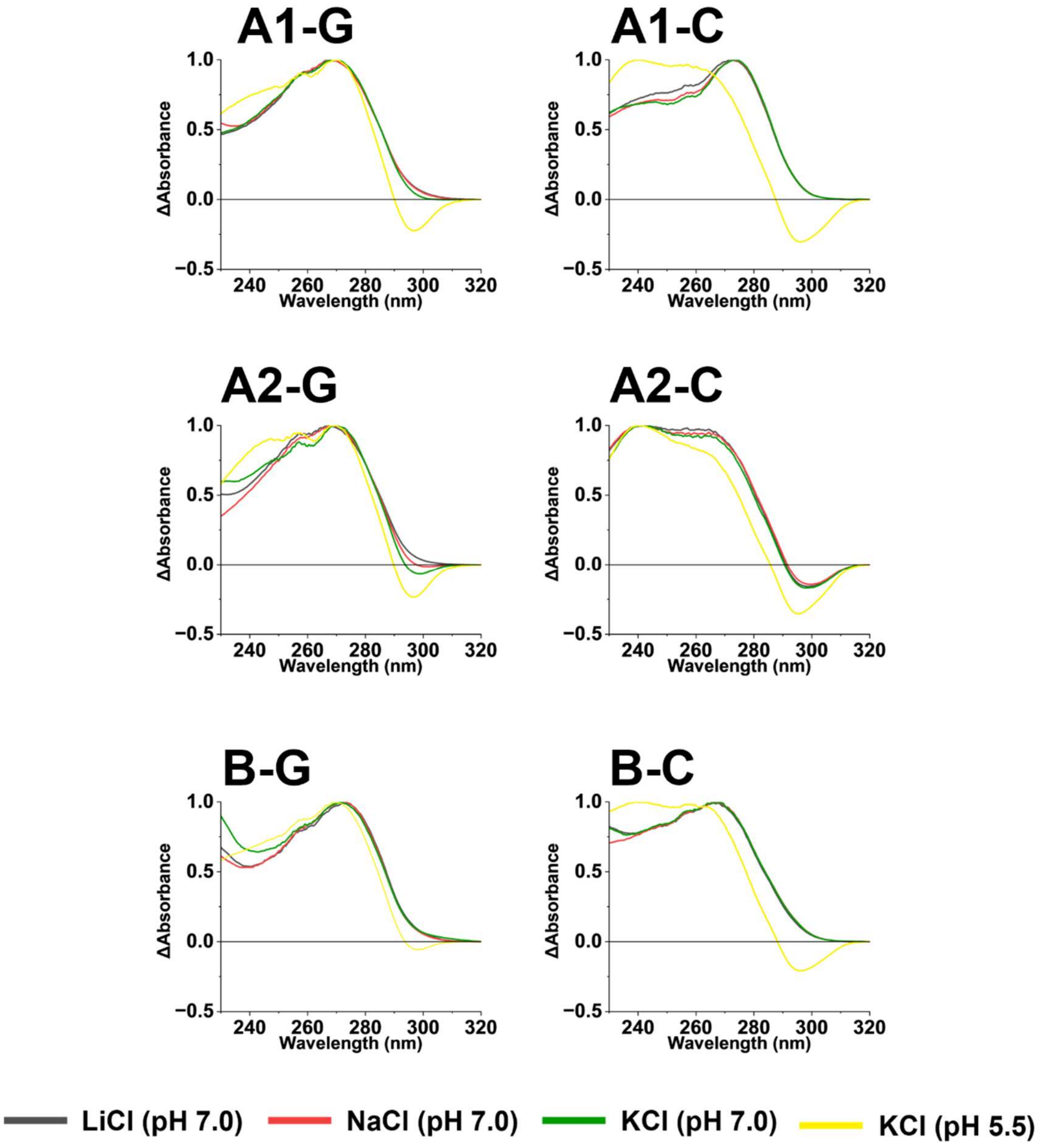


**Supplementary Fig. 4.** Thermal difference spectra of the *cyp51* sequences. DNA oligonucleotides (5 μM) were annealed in 10 mM sodium cacodylate buffer containing 100 mM LiCl, NaCl, KCl (pH 7.0), or KCl (pH 5.5) as indicated.

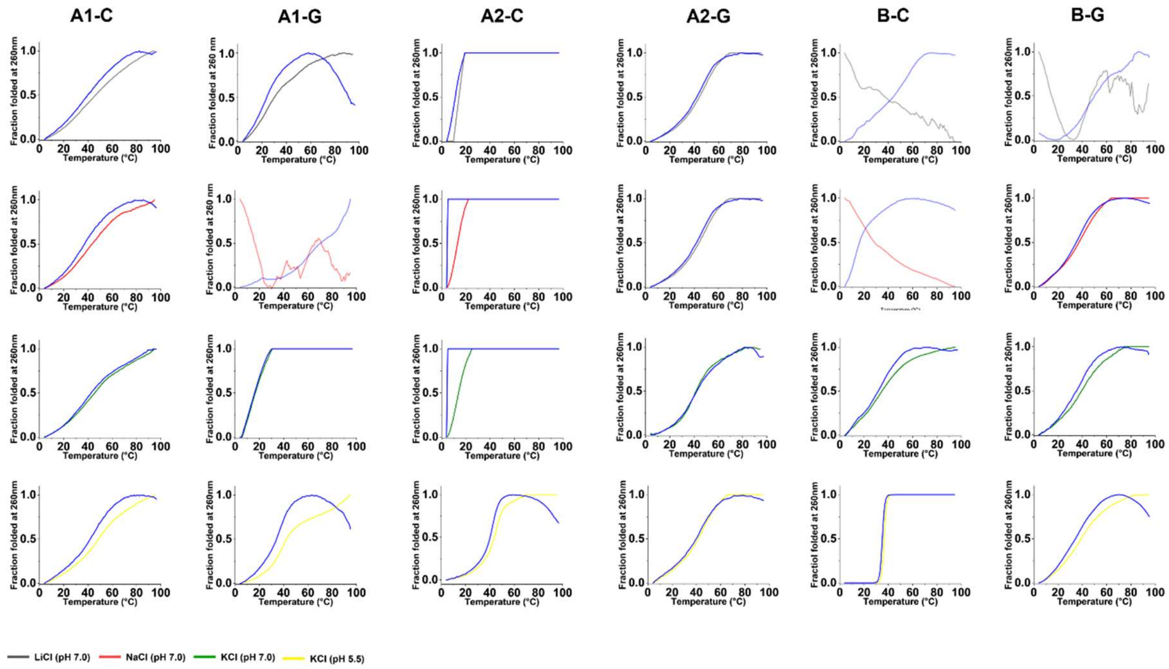


**Supplementary Fig. 5.** UV melting and annealing profile of 5 μM of the DNA oligonucleotides A1-C, A1-G, A2-C, A2-G, B-C, and B-G at 260 nm in 10 mM sodium cacodylate buffer containing 10 mM LiCl, NaCl, KCl (all at pH 7.0) and KCl (at pH 5.5).

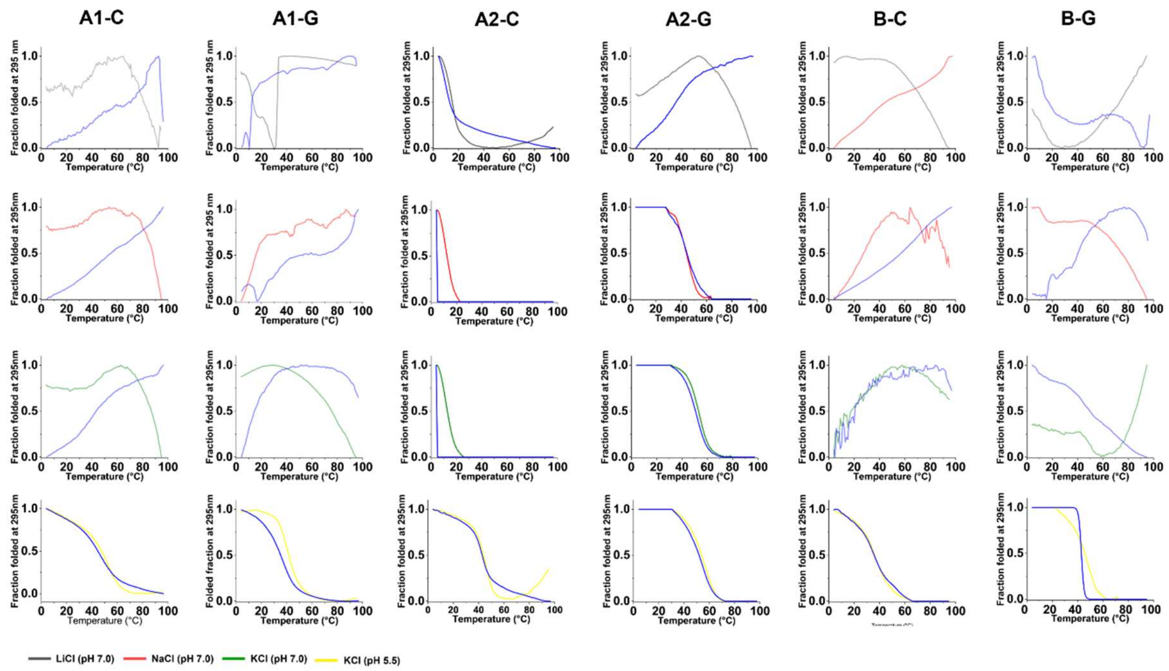


**Supplementary Fig. 6**. UV melting and annealing profile of 5 μM of the DNA oligonucleotides A1-C, A1-G, A2-C, A2-G, B-C, and B-G at 295 nm in 10 mM sodium cacodylate buffer containing 10 mM LiCl, NaCl, KCl (all at pH 7.0) and KCl (at pH 5.5).

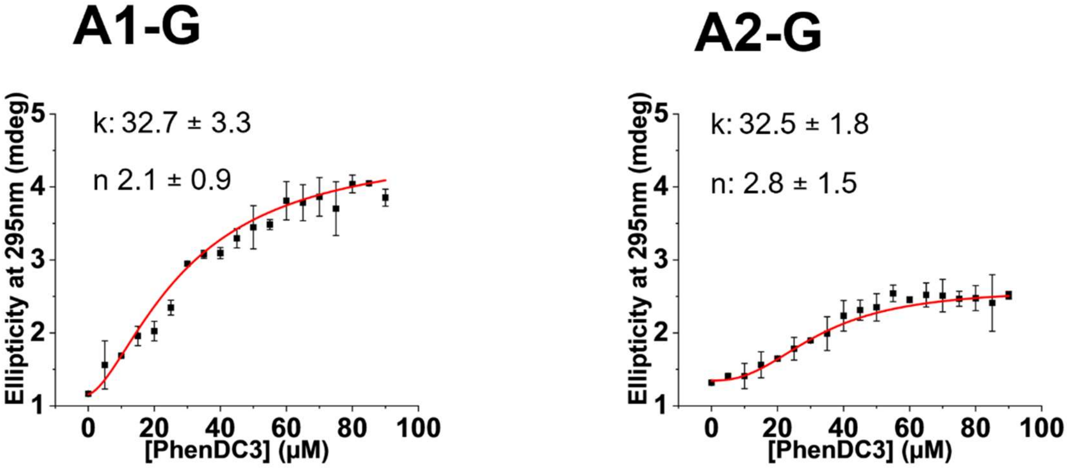


**Supplementary Fig. 7.** Plot of ellipticity of the experimental repeats in Fig. 1d at 295nm against PhenDC3 concentration and the corresponding Hill-1 fitting.

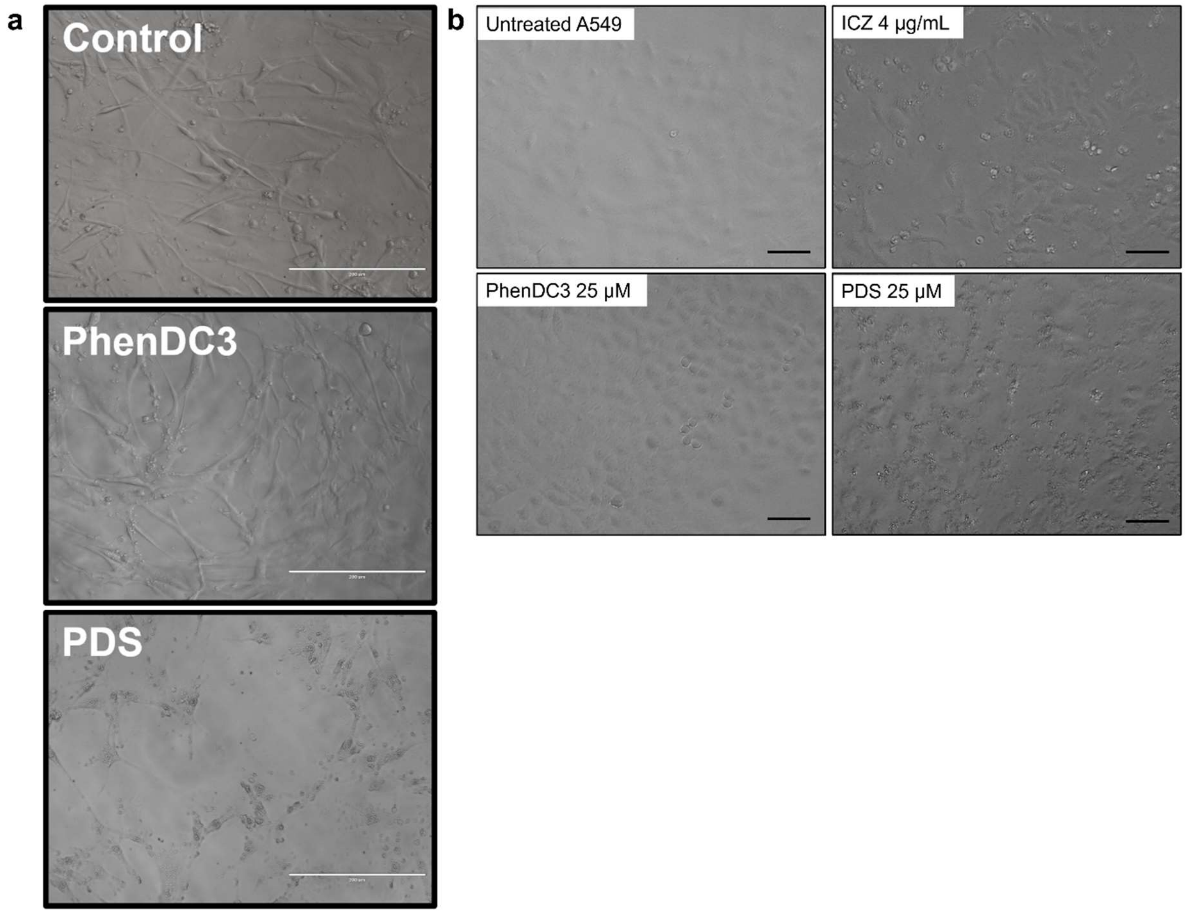


**Supplementary Fig. 8.** Representative light microscopy images of (**a**) primary human vascular smooth muscle cells treated with 25 µM PhenDC3 or PDS for 24 h. Scale bars represent 200 µm. Representative light microscopy images of (**b**) human A549 lung cells treated with 25 µM PhenDC3, 25 µM PDS, or 4 µg/mL itraconazole (ICZ) for 24 h. Scale bars represent 100 µm.

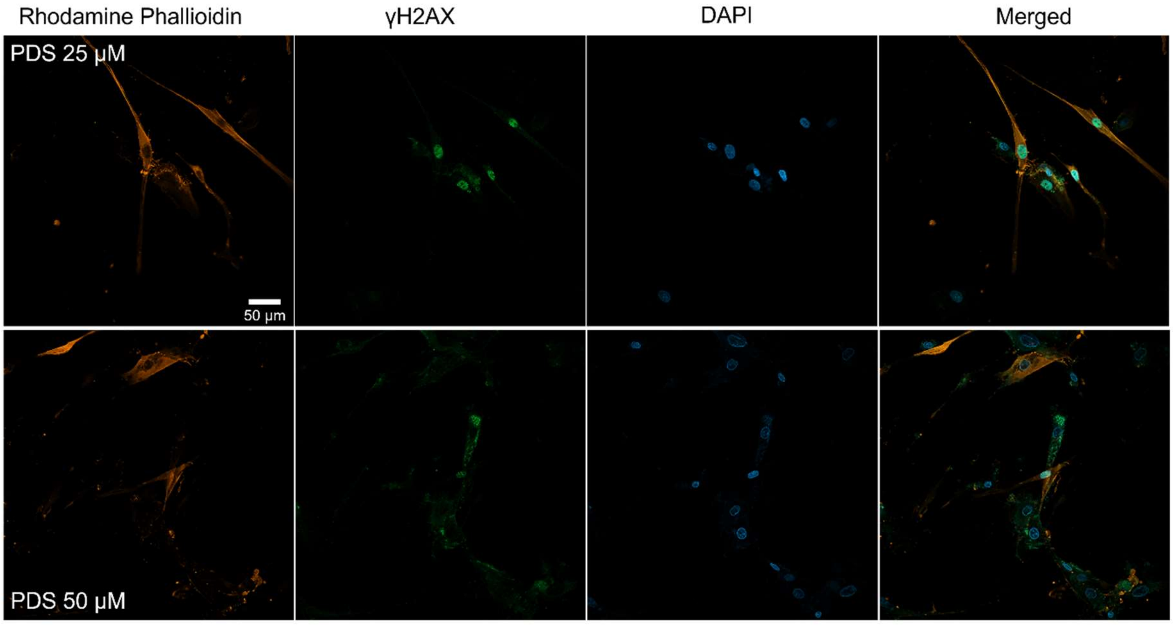


**Supplementary Fig. 9.** Representative confocal microscopy images of primary human vascular smooth muscle cells treated with 25 µM or 50 µM PDS for 24 h. Scale bars represent 50 µm. Cells were incubated with rhodamine phalloidin, a γH2AX antibody, or DAPI to visualise F-actin, DNA damage, or the nucleus, respectively. Images were obtained using a Zeiss LSM 980 laser scanning confocal microscope equipped with an Airyscan 2 detector using the 20x objective. Images were analysed using Fiji 1.53t.
